# Mitf is a Schwann Cell Sensor of Axonal Integrity that Drives Nerve Repair

**DOI:** 10.1101/2022.09.25.509350

**Authors:** Lydia Daboussi, Giancarlo Costaguta, Miriam Gullo, Nicole Jasinski, Veronica Pessino, Brendan O’Leary, Karen Lettieri, Shawn Driscoll, Samuel L. Pfaff

**Affiliations:** Gene Expression Laboratory, Salk Institute for Biological Studies, 10010 North Torrey Pines, La Jolla, CA 92037, USA

## Abstract

Schwann cells respond to acute axon damage by transiently transdifferentiating into specialized repair cells that restore sensorimotor function. However, the molecular systems controlling repair cell formation and function are not well defined and consequently it is unclear whether this form of cellular plasticity has a role in peripheral neuropathies. Here we identify Mitf as a transcriptional sensor of axon damage under the control of Nrg-ErbB-PI3K-PI5K-mTorc2 signaling. Mitf regulates a core transcriptional program for generating functional repair Schwann cells following injury and during peripheral neuropathies caused by CMT4J and CMT4D. In the absence of *Mitf*, core genes for epithelial-to-mesenchymal transition, metabolism and dedifferentiation are misexpressed and nerve repair is disrupted. Taken together, our findings demonstrate that Schwann cells monitor axonal health using a phosphoinositide signaling system that controls Mitf, which is critical for activating cellular plasticity and counteracting neural disease.

**Highlights:**

- Mitf-induced Schwann cell plasticity is triggered by peripheral neuropathy.
- Nrg-ErbB signaling activates Mitf via cytoplasmic-to-nuclear translocation.
- Mitf restores sensorimotor function following axonal breakdown.
- Mitf regulates a core repair program across both injury and neurodegeneration.

## Introduction

In response to injury many tissues undergo adaptive cellular reprogramming to generate specialized repair cells for wound healing ^1, 2^. Intuitively this trans-differentiation process could be considered developmental regression involving factors from embryogenesis, however repair cell generation appears to engage separate pathways dedicated uniquely for injury ^1^ . While the central nervous system has a limited regenerative capacity - the peripheral nervous system possesses an astonishing capability for self-repair. Axonal damage caused by trauma induces mature Schwann cells (SCs), a type of glia within peripheral nerves, to transiently convert into repair cells that restore nerve function ^3, 4^. While it is established that SC plasticity is fundamental for healing and repair, the molecular sensor(s) within SCs that detect axonal damage and safely program mature SCs into repair cells with temporal and spatial precision are not well characterized.

Developmental studies have revealed many cellular steps and molecular factors involved in neural crest development into melanoblasts and SCs (Figure 1A) ^5^. Mature SCs ensheathe peripheral axons as either myelinating or non-myelinating (Remak) cells ^6^. Upon peripheral nerve injury, loss of axonal contact causes both myelinating and Remak SCs to be reprogrammed into repair SCs (Figure 1A) ^3^. The transformation of mature but plastic SCs into repair cells requires reprogramming of the transcriptome leading to expression of genes associated with stemness (i.e. genes associated with self-renewal), in addition to factors controlling an epithelial-to-mesenchymal (EMT) conversion thereby promoting tissue remodeling and injury repair ^7–10^. These repair cells perform specialized functions not carried out by either mature SCs or SC precursors, including the clearance of axonal debris, activation of innate immune responses, and formation of Büngner bands that serve as a substrate for the regrowth of axons to peripheral targets ^11^. Some of the factors controlling repair cell function have begun to be identified including c-Jun necessary for myelin clearance and Stat3 involved in maintaining repair cell survival ^12–17^.

**Figure 1.**
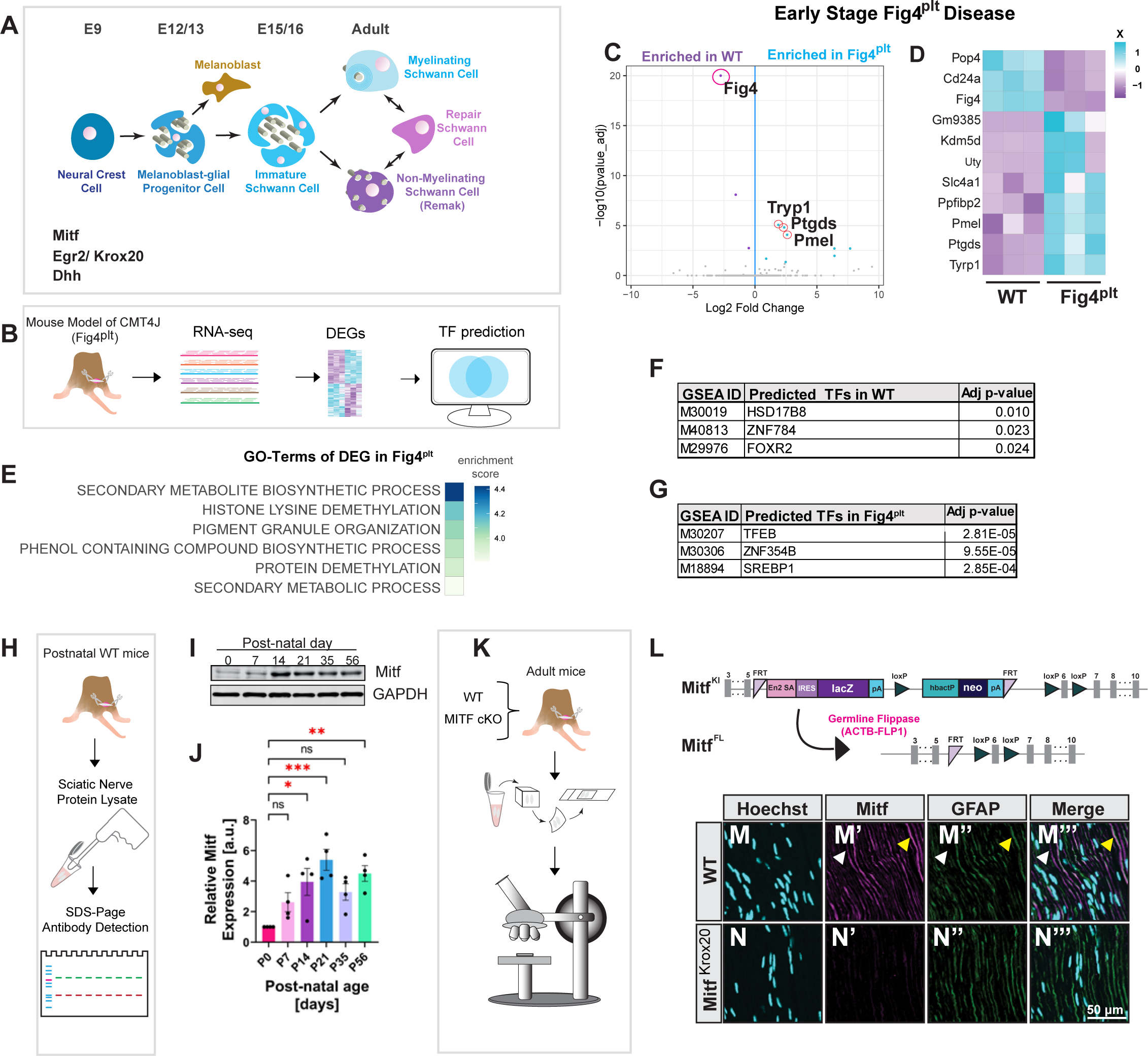
Network Analysis Links Mit/Tfe to Schwann cell response in CMT4J. **(A)** Diagram of Schwann cell development and a subset of molecular markers that delineate each stage. **(B)** Workflow for the identification of differentially expressed genes and transcription factor motif enrichment (GSEA). **(C)** Volcano plot showing log fold change versus adjusted p-value of sciatic nerves of Fig4^plt^ at early-stage disease (P4). Each dot represents a gene. Blue and purple dots are differentially expressed. N=3, each sample is pooled from both sciatic nerves of 3 animals. **(D)** Heat map of expression levels (Z-score) of differentially expressed genes from **(C)**. Each row corresponds to a gene. **(E)** Biological Process (BP) GO-Terms from differentially expressed genes (DEG). **(F-G)** Gene Set Enrichment Analysis (GSEA) of putative transcription factor binding motifs in either WT **(F)** or Fig4^plt^ (**G)**. **(H)** Sciatic nerve protein lysate schematic of postnatal WT mice. **(I)** Protein lysates of sciatic nerves from WT (ICR) animals analyzed at P0, 7, 14, 21, 35, and 56 by SDS-page and immunoblotting. **(J)** Quantification of **(H)** shows Mitf protein expression normalized against GAPDH and measure relative to the expression at P0 (n=4). Ordinary One-way ANOVA with Dunnett’s multiple comparison test. **(K)** Sciatic nerve immunofluorescence schematic. **(L)** Mitf knock-in construct and genetic strategy to convert the Mitf knock-in allele (Mitf^KI^) to the Mitf floxed allele (Mitf^FL^). **(M-N’’’)** Longitudinal sections of adult sciatic nerves (12 wks) were co-immunostained for Mitf and GFAP and costained with Hoechst in either **(M)** or Mitf^Krox^^20^ **(N)**. Yellow Arrowhead shows Mitf colocalization with GFAP. White arrowhead shows Mitf staining without GFAP.

Beyond physical injuries, nerve function is also compromised in hereditary peripheral neuropathies including many forms of Charcot-Marie-Tooth (CMT) disease. Genetic studies have identified >10 recessive genes for CMT Type IV (CMT4) alone, affecting axon myelination, SC-axon interactions and ultimately neuronal survival. Thus, CMT4 patients often experience muscle weakness and a diminished touch and thermal sensitivity in their distal limbs ^18^. Candidate factors revealed from human genetics include the phosphoinositide 5-phosphatase encoded by the *Fig4* gene mutated in CMT4J, the phosphatidylinositol 3-kinase (PI3K) inhibitor *Ndrg1* mutated in CMT4D^19, 20^, the Rab11 effector *SH3TC2* mutated in CMT4C ^21, 22^, and the PI(3)P phosphatase *MTMR2* mutated in CMT4B ^23–25^. Taken together, the genes linked to CMT4 suggest that second messenger signaling via lipid phosphoinositides may control how SCs respond to extracellular cues ^26, 27^. While it is clear that plasticity is critical for repair following injury, the role for plasticity in neural disease remains enigmatic.

We investigated whether repair SCs are generated and modify disease progression in different types of CMT4 mouse models. To identify how the contact between axons and SCs is sensed we characterized the transcriptome of nerves from CMT4J (*Fig4^plt^*) mice in which the close apposition of SCs to motor and sensory axons is disrupted. Bioinformatics predicted that target genes of the Mit/Tfe family of bHLH transcription factors were altered in CMT4J, suggesting Mit/Tfe factors are linked to the disease. Mitf, a member of this family, was genetically identified eight decades ago from selection of naturally-occurring *Mitf*-mutant mice that displayed altered pigmentation and coat color ^28^. The subsequent cloning and characterization of Mitf has revealed this transcription factor is necessary for the embryonic development of neural crest-derived cell types, including melanocytes (Figure 1A) ^29–31^. We discovered that Mitf is also expressed by mature SCs, and that disrupting axon-SC interactions by nerve injury induces the transcription factor to translocate from the cytoplasm to the nucleus. We found that axon damage triggers Mitf nuclear localization by disrupting Nrg1-ErbB-PI3K-PI5K-mTorc2 signaling. Furthermore, we discovered that Schwann cells induce a shared transcriptomic response that mediates repair in both injury and disease. Mitf regulates this core transcriptional program in injury and disease-derived repair cells controlling EMT, metabolic and trophic factors, and SC dedifferentiation. Consequently, when *Mitf* is mutated nerve repair is disrupted and peripheral neuropathies such as CMT4D are more severe. Our findings indicate that Mitf controls a core network of genes linked to SC plasticity and repair cell function that modifies how peripheral neuropathies manifest.

## Results

### Network Analysis Links Mit/Tfe bHLH Family to CMT4J

Mutations in the lipid phosphatase encoded by the *Fig4* gene cause a demyelinating form of Charcot-Marie-Tooth, termed CMT4J, that leads to muscle weakness in humans. To identify Schwann cell responses to CMT4J disease we sequenced sciatic nerve RNA from the “*pale tremor*” mouse line which carries a transposon insertion that disrupts *Fig4* (Fig4^plt^) ^32, 33^. Comparison of the transcriptome of Fig4^plt^ to wild type (WT) mice during mid-stage disease revealed 485 genes were upregulated and 473 genes were downregulated. (Figure S1A-B). GO-Term (Gene Ontology) analysis identified several pathways related to mature SC function enriched in the controls (Figure S1C). Conversely, genes associated with mature SCs were downregulated in Fig4^plt^ mice, whereas injury-response and wound healing pathways were upregulated (Figure S1D). These data suggest that CMT4J peripheral neuropathy triggers a conversion of SCs from a mature myelinating state to a repair/wound healing state. We used Gene Set Enrichment Analysis (GSEA) to identify Transcription Factor (TF) binding motifs enriched in the promoter regions of differentially expressed genes in Fig4^plt^ and found motif-enrichment of 93 TFs in the control and 68 in the Fig4^plt^ mutant. Among the candidates, we detected AP-1 motif enrichment in the differentially expressed transcriptome of *Fig4^plt^*, suggesting c-Jun may function during CMT4J (Figure S1E-F)^12^. These data underscore the dramatic transformation of the peripheral nerve during mid-stage disease progression of *Fig4^plt^*.

To understand the genes and pathways linked specifically to the early phase of the CMT4J disease-response we sequenced sciatic nerve RNA from Fig4^plt^ mice at early-stage disease (P4) (Figure 1B-D). RNA-sequencing revealed that Fig4 RNA is enriched in the WT nerve at this early age. GO-Term analysis indicated that ‘Metabolic’, ‘Biosynthetic Process’ and ‘Pigment Granule Organization’ pathways were upregulated in the Fig4^plt^ mutant compared to WT (Figure 1E). This was unexpected because post-natal peripheral nerves do not contain melanin producing cell-types, this implies that common cellular pathways that result in Fig4^plt^ hypopigmentation are also disrupted in Schwann cells. Furthermore, at early-stage disease GSEA revealed motif enrichment of 11 TFs in Fig4^plt^, and 4 TFs in WT (Figure 1F-G). Tfeb-family bHLH transcription factors (Tfeb, Tfe3, Tfec, Mitf) showed the highest enrichment in the differentially expressed genes in sciatic nerves from Fig4^plt^ mutants (Figure 1G). Interestingly, TFEB promoter elements were also enriched in the differential gene expression at later disease time points (Figure S1F), indicating one or more members of this gene family may regulate several stages of disease response in CMT4J.

### Mitf Is Enriched In Schwann Cells

We investigated the role that Mitf, a Mit/Tfe family member, plays in SCs because: (1) there are pigmentation defects in Fig4^plt^ mutants and Mitf is a master regulator of melanocyte development and (2) Mitf has been shown to directly regulate several of the differentially expressed genes at P4 including Pmel, Ptgds and Tryp1 ^34, 35^ (Figure 1C, D). Previous studies have shown that Mitf is transiently expressed in the neural crest of the developing embryo and is down-regulated as the melanoblast-glia progenitor cells differentiate into SCs ^31, 36^ . However, the role of Mitf in SCs is complicated by the observation that this factor is necessary for the generation of some neural crest derived cell-types such as RPE (retinal pigment epithelial) cells and melanocytes in mice^30^. This leaves open the question as to whether there is a specific role for Mitf in Schwann cells or only an early function in the neural crest derived precursors of these glia. To address this, we first generated Mitf primary antibodies and conducted a time-course of protein expression in the sciatic nerve. Low levels of Mitf were detected in the peripheral nerve at birth followed by an increase at P14 that is sustained through adulthood (Figure 1H-J). Thus Mitf has a biphasic expression during SC development - first in neural crest, and then in mature SCs (Figure 1A). In the adult nerve, our analysis revealed Mitf is expressed by both GFAP**^+^** cells (nonmyelinating; Remak) and GFAP**^-^** (myelinating SCs) (Figure 1K-N’’’). While these data reveal that Mitf is expressed in mature SCs, we unexpectedly found that it is enriched in the cytoplasm rather than nuclei (Figure 1M-M’’’).

### Generation of Conditional *Mitf* Alleles

Constitutive mutations of *Mitf* have revealed its early role in neural crest development but this obscures the possible function in mature SCs^37^ . Therefore, we generated a conditional *Mitf* allele. First, a *Mitf* knock-in mouse (Mitf^KI^) was made using “knockout-first” targeting constructs (Figure 1L). Homozygous animals for this knock-in allele (Mitf^KI/KI^) lack Mitf protein expression, and are therefore equivalent to a constitutive knockout (Figure S1I). Consistent with previous *Mitf* loss-of-function mutations, Mitf^KI/KI^ mice were albino, had a reduced startle response, and had smaller eyes (microphthalmia) compared to littermate controls ^38, 39^(Figure S1G-I). Despite these changes Mitf^KI/KI^ mice did not exhibit signs of peripheral nerve dysfunction and displayed normal locomotor activity consistent with previous descriptions of *Mitf* mutants ^40^.

Next, we converted the Mitf^KI^ allele to a conditional-flox allele (Mitf^fl^), encoding wild-type protein, by crossing to a mouse line with germline Flp expression (Figure 1L). To test the conditional nature of the Mitf^fl/fl^ allele we used CDX2:CRE to selectively delete the *Mitf* gene from the caudal region of developing embryos (Mitf^CDX2^; Figure S1J-K). As expected, Mitf^CDX2^ animals display caudal-selective albinism due to deletion of *Mitf* during caudal melanocyte development. Deletion of *Mitf* from the caudal embryo did not overtly affect peripheral nerve function in adult animals, consistent with null animals (Figure S1H) ^29^.

### Mitf is Dispensable for Schwann Cell Development

While hypopigmentation defects arise in mice and humans with *Mitf* mutations, peripheral nerve-related phenotypes have not been described ^29^. However, this did not preclude a subtle role for Mitf in SCs exposed to specific physiological conditions. Crossing the Mitf^fl/fl^ line to the pan-SC line Krox20:CRE (Mitf^Krox20^) selectively and efficiently ablated Mitf expression in peripheral nerve SCs (Figure 1L-N’’’). We examined the sciatic nerves of Mitf^Krox20^ and WT mice at P7 and 3 months of age using transmission electron microscopy (TEM). At both time points, *Mitf* mutants exhibited normal myelin thickness (g-ratio), axon diameter, Remak bundles and extra cellular matrix organization (Figure 2A-E, S2A-E). Mitf^Krox20^ animals did not show alterations in grip strength at 3, 6, or 12 months of age, or rotorod motor coordination defects in adults (Figure 2F-G; S2F-J).

**Figure 2.**
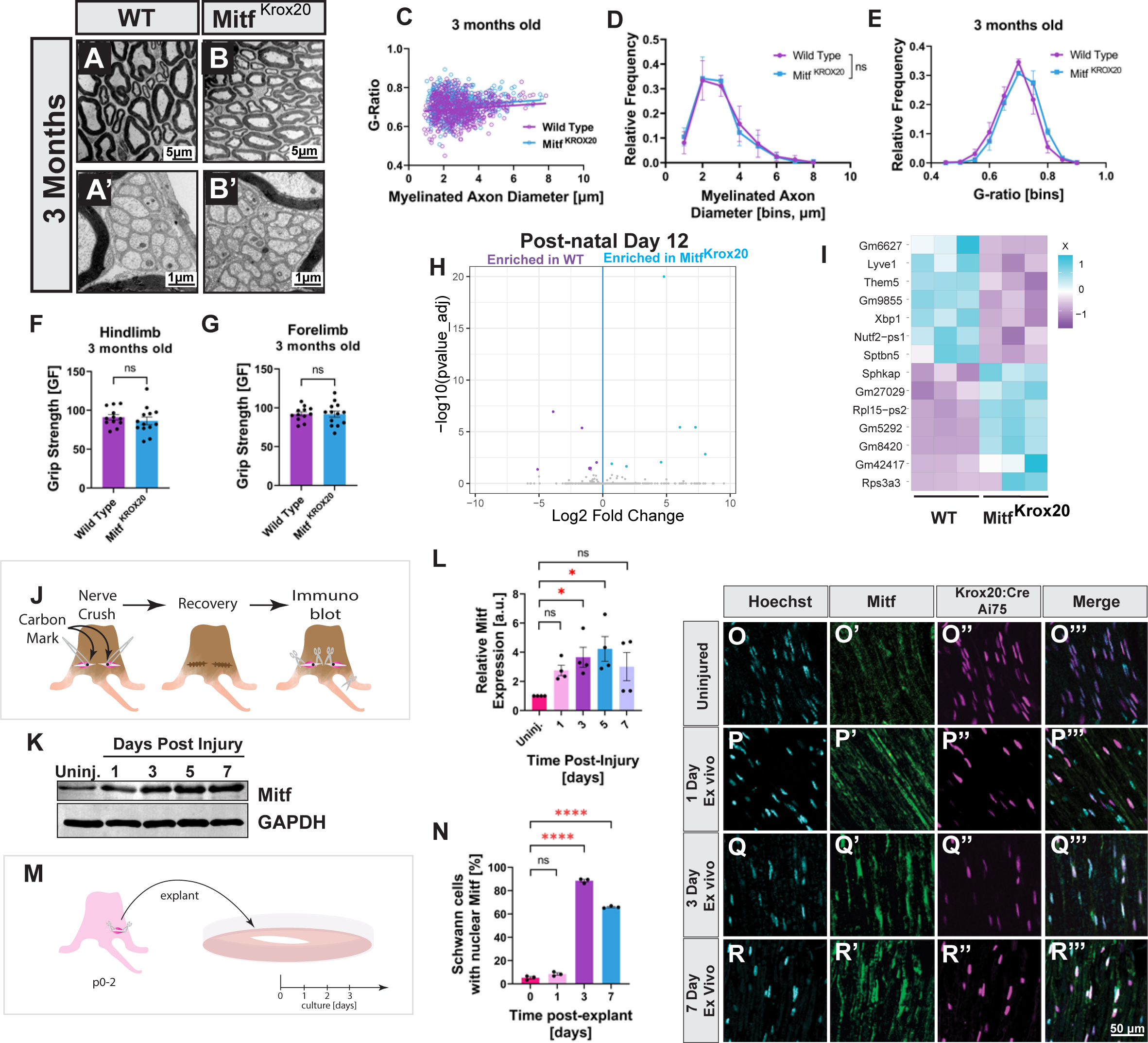
Mitf is Dispensable for Development and Upregulated in Schwann cells during Wallerian Degeneration. **(A-B’)** TEM of sciatic nerve cross-sections at 3 months show myelinating Schwann cells in either WT **(A)** or Mitf^Krox^^20^ **(B)** and non-myelinating Schwann cells (A’, B’). **(C)** G-ratio plotted against axon diameter of 3 months old WT (purple) and Mitf^Krox^^20^ (blue) (n=3, at least 200 axons counted per sample). Lines represent simple linear regression. **(D)** Frequency of axon diameter in either WT or Mitf^Krox^^20^. Multiple unpaired t-Test **(E)** G-ratio frequency of WT and Mitf^Krox20^ (N=3, > 200 axons counted per sample). Multiple unpaired t-Test **(F-G)** Hindlimb **(F)** and forelimb **(G)** grip strength at 3 months old for WT (N=9) and Mitf^Krox20^ (n=13). Unpaired t-Test. **(H)** Volcano plot showing log fold change versus adjusted p-value of sciatic nerve of control or Mitf^Krox20^ at P12. Each dot represents a gene, blue and purple dots are differentially expressed. N=3, each sample is pooled from both sciatic nerves of 3 animals. **(I)** Heat map of relative expression levels (Z-score) of differentially expressed genes from **(H)**. Each row corresponds to a gene. **(J)** Diagram of the nerve crush. Nerves were crushed under the sciatic notch and marked with carbon. Animals allowed to recover for 1, 3, 5, or 7 days. **(K)** Protein lysates of sciatic nerves from animals 1, 3, 5, 7 dpi were analyzed by SDS-page and immunoblotted for Mitf and GAPDH. Each lane was loaded with 5 µg of protein. **(L)** Quantification of immunoblots from **(K)**. Mitf expression was normalized in each experiment relative to the uninjured nerve (lane 1); N=4. Ordinary One-way ANOVA with Dunnett’s multiple comparison test. **(M)** Diagram of sciatic nerve explant culture. Nerves were cultured ex vivo for 1, 3, or 7 days thereby inducing Wallerian degeneration. **(N)** Quantification of the number of Schwann cell nuclei that show staining for Mitf. Schwann cell nuclei are specifically labelled with an nls-Td-tomato reporter (Ai75) driven by Krox20-CRE (n=3). Ordinary One-way ANOVA with Dunnett’s multiple comparison test. **(O-R’’’)** Genetically labelled Schwann cells in the sciatic nerve (Krox20-CRE; Ai75) were either acutely chemically fixed **(O-O’’’)** or cultured ex-vivo for 1, 3, or 7 days **(P-R’’’)** and then chemically fixed.

To identify molecular changes that might be more subtle than behavioral alterations we performed RNA-sequencing on sciatic nerves from Mitf^Krox20^ and littermate controls. We found 15 differentially expressed transcripts, 9 of which represented protein coding mRNAs (Figure 2H-I). We independently confirmed these findings by deleting Mitf from SCs using DHH:CRE and identified 13 differentially expressed genes (Figure S2K-L). However, none of the transcripts from the outbred, mixed genetic backgrounds comprising Mitf^DHH^ overlapped with Mitf^Krox20^, suggesting that the loss of Mitf in mature SCs leads to few, if any, detectable RNA expression changes. Taken together, the similarities in molecular, cellular and functional characteristics between WT and conditional *Mitf* mutant mice suggest that this transcription factor is dispensable for the normal development and function of mature SCs.

### Nerve Injury Induces Nuclear Mitf Translocation

Although Mitf is expressed in SCs, it is localized to the cytoplasm and the absence of phenotypes in *Mitf* mutants suggests it has no function outside the nucleus. This prompted us to test whether Mitf functions in SCs under non-homeostatic conditions, such as those induced by nerve injury. First, we examined whether Mitf localization changes following nerve transection using an ex-vivo preparation that prevents axonal sprouting and lymphocyte infiltration. Similar to the uninjured nerve at day 1 Mitf was localized to the cytoplasm; whereas by day 3, 89% of SC nuclei had enriched staining for Mitf (Figure 2M-R’’’). Next, we examined Mitf protein levels in vivo following nerve crush injury. With similar kinetics to the nuclear relocalization of Mitf, at day 3 we observed a 3.6 fold increase of overall protein levels (Figure 2J-L). Interestingly these injury-induced increases in Mitf protein were not associated with increased RNA, indicating that Mitf protein levels are post-transcriptionally regulated (Figure S2M). These results indicate that axon integrity dynamically influences the protein concentration and localization of Mitf likely through a post-translational mechanism.

### Mitf is Required for Peripheral Nerve Regeneration

The increase in Mitf protein and nuclear localization in SCs following nerve injury prompted us to investigate whether it has a specific role in nerve regeneration. To identify the time points at which loss of Mitf might impact nerve recovery we conducted a histological time course at 3, 5, 21, 42 and 90 days after peripheral nerve crush (Figure 3A). At 3 and 5 dpi, we found nerves from Mitf^Krox20^ mice undergo faster axonal degeneration (i.e. Wallerian degeneration) as evidenced by fewer myelinated axons (Figure 3B-F, S3A-D). These findings were striking because the rapid degeneration observed in *Mitf* mutants is in direct contrast to the slow Wallerian degeneration observed in WldS and *c-Jun* cKO mouse models ^12, 41^. At 21dpi, SCs exhibited a diminished small caliber axon-Schwann cell relationship (Figure S3E-F’). These early perturbations result in a series of downstream consequences. At 42 dpi, a striking collection of SC defects were observed in Mitf^Krox20^ animals and included: ectopic myelination of small caliber axons, supernumerary wrapping of small unmyelinated axons, SC wrapping of collagen, enduring Büngner SCs and axons misdirected through the perineurium (Figure 3I-K, S3G-G’’’). At 90 dpi in *Mitf* mutants, SCs wrapped around collagen fibrils, and had lingering SC processes present throughout the extra cellular matrix (ECM). While the development of myelin is normal in *Mitf* mutants, remyelination following injury was altered leading to thicker myelin (smaller G-ratio) and a decrease in myelinated axon caliber (Figure 3L-P). Taken together, these data suggest that Mitf function is required early in the SC transdifferentiation process and that its loss permanently impacts Schwann cell-axon physical interactions necessary for proper nerve repair.

**Figure 3.**
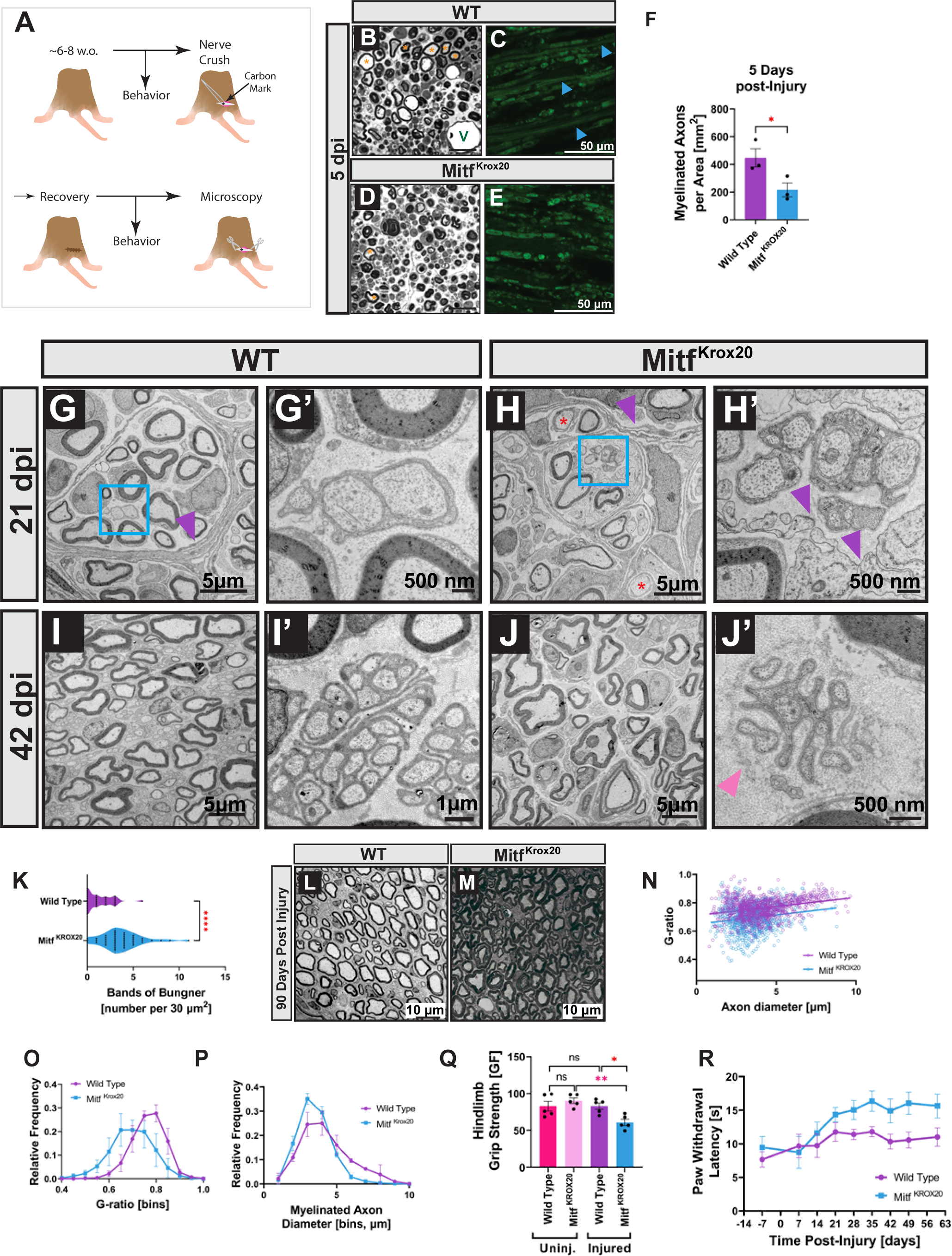
Mitf Is Required For The Normal Repair Of The Nerve After Injury. **(A)** Diagram of nerve crush in adult animals 6-8 weeks old. **(B,D)** Toluidine blue-stained semi-thin sections of WT or Mitf^Krox20^ 5 days post injury, quantified in **(D)**. (Orange Asterisk =myelinated axons, V=Vessel, White arrowhead= intact myelinated axon). **(C,E)** Fluoromyelin stained longitudinal sections of sciatic nerve 5 days post injury. (Blue arrowheads indicate uninterrupted myelin staining). **(F)** Quantification of the number of intact myelinated axons in either WT or Mitf^Krox20^ adult mice 5 days post injury (n=3). Unpaired t-Test. **(G-H’)** Representative TEM cross-sections of the sciatic nerve bridge 21 days after transection. Showing normal (purple arrow, **G**) and abnormal perineurial organization (purple arrow, **H**) and fibroblast infiltration (purple arrow, **H’**). **(I-J’)** TEM images of a sciatic nerve cross section 5 mm below sciatic nerve crush show either normal Schwann cells wrapping of the control (I,-I’) or persistence of Büngner Schwann cells and aberrant axon-SC interactions in Mitf^Krox20^ (pink arrowhead, **J-J’**). **(K)** Quantification of Bϋngner cells present at 42 days post sciatic nerve crush 5 mm distal from the site of the nerve injury in TEM cross sections. N=3 control; N=4 mutant, >5 fields per animal. Unpaired t-Test. **(L,M)** Representative TEM 3 months after sciatic nerve injury of either WT **(L)** or Mitf^cKO^ **(M)**, 5 mm from the site of injury. Scale Bar = 10 µM. **(N)** G-ratio versus axon diameter of 3 months after injury, 5 mm distal from the nerve crush in control (purple) and Mitf^Krox20^(blue) (n=3, >150 axons per animal). Lines represent simple linear regression. **(O)** G-ratio frequency of WT and Mitf^Krox20^ (N=3, > 200 axons counted per sample). Multiple unpaired t-Test **(P)** Quantification of the frequency of myelinated axon diameter in control and Mitf^Krox20^ sciatic nerve 3 months after injury, 5 mm from the site of injury (n=3, >150 axons per animal). Multiple unpaired t-Test. **(Q)** Hindlimb grip strength of WT and Mitf^Krox20^ 60 days after sciatic nerve pinch of the injured hind limb or the contralateral uninjured hindlimb (n=5). Grams of Force (GF). Ordinary One-way ANOVA with Šidák’s multiple comparison test. **(R)** Hargreaves assay measuring paw withdrawal latency over time of WT and Mitf^Krox20^ animals 7 days before injury through 60 days after sciatic nerve crush (n=5-8). Mixed effects ANOVA analysis with Geisser-Greenhouse’s correction: column factor (WT v Mitf^Krox20^) p = 0.0118 (*), F (1, 14) = 8.364; row factor (time) p = 0.0016 (**) F (4.110, 41.61) = 5.189; column x row factor p = 0.3392 (ns), F (8, 81) = 1.151.

After transection injuries, which create a gap in the nerve, unlike a crush, repair SCs take on increased mesenchymal characteristics as they invade the gap between the distal and proximal stump and guide axons through the nerve bridge ^9^. Interestingly, an additional layer of nerve disorganization was revealed after nerve transection in the *Mitf* mutants. At 21 days after sciatic nerve transection we found disorganized, patchy perineurial fibroblast organization, the existence of an inappropriately infiltrating fibroblast cell-type, and unmyelinated axons (Figure 3G-H’). After injury, repair SCs facilitate nerve reconstruction in part through signals to connective tissue^42^. Our results indicate that this process is disrupted in *Mitf* mutants.

We investigated whether the cellular changes we observe in *Mitf* mutants manifested as behavioral alterations in sensorimotor function following nerve crush. Prior to sciatic nerve injury both WT and Mitf^Krox20^ mice responded with similar latencies to noxious heat on their hind paws. At 28 through 60 dpi, Mitf^Krox20^ animals had significantly longer latencies to this stimulus compared to controls, indicating the recovery of nociceptive sensory function was impaired (Figure 3R). Next, we tested hindlimb grip strength. Prior to injury WT and Mitf^Krox20^ mice displayed similar grip strength, whereas at 60 dpi the grip strength recovery of injured *Mitf* mutants was significantly lower than the controls (Figure 3Q). These deficits in recovery of sensorimotor function following injury indicate that Mitf is required for effective repair of damaged nerves in vivo.

### Mitf Regulates a Dynamic Gene Network in Nerves Undergoing Repair

To investigate the molecular basis by which Mitf mediates repair we conducted a whole-genome RNA sequencing time-course following sciatic nerve transection at 3, 7, and 14 dpi (Figure 4A). After injury, mature SCs transition into repair SCs by undergoing EMT and acquiring a phagocytic phenotype^8, 9^ (Figure 4B). EMT is mediated by the downregulation of adhesion pathways involving WNT and TGF-β signaling components ^8, 9, 43^. At 3 dpi WT SCs differentially expressed 74 genes relative to *Mitf* mutants (34 upregulated, 39 downregulated) (Figure 4C). Then, at 7 dpi, we detected 345 differentially expressed transcripts (244 upregulated and 101 downregulated) in WT nerves relative to *Mitf* mutants (Figure 4D). Although Mitf has been linked to lysosomal biogenesis and autophagy neither of these pathways were significantly affected^44–47^ (Figure S4E-F). Rather, Mitf^Krox20^ animals exhibited dysregulation of genes linked to: cellular adhesion and extracellular matrix organization, metabolism and glucose/lactate production, signaling molecules (including TGF-β and WNT signaling components), and elevated mature SC markers (Figure 4D, F; S4A- B). Taken together, these data suggest that alterations in *Mitf* mutants block proper SC differentiation leading to altered EMT and energy metabolism, early processes essential for the function of repair SCs.

**Figure 4.**
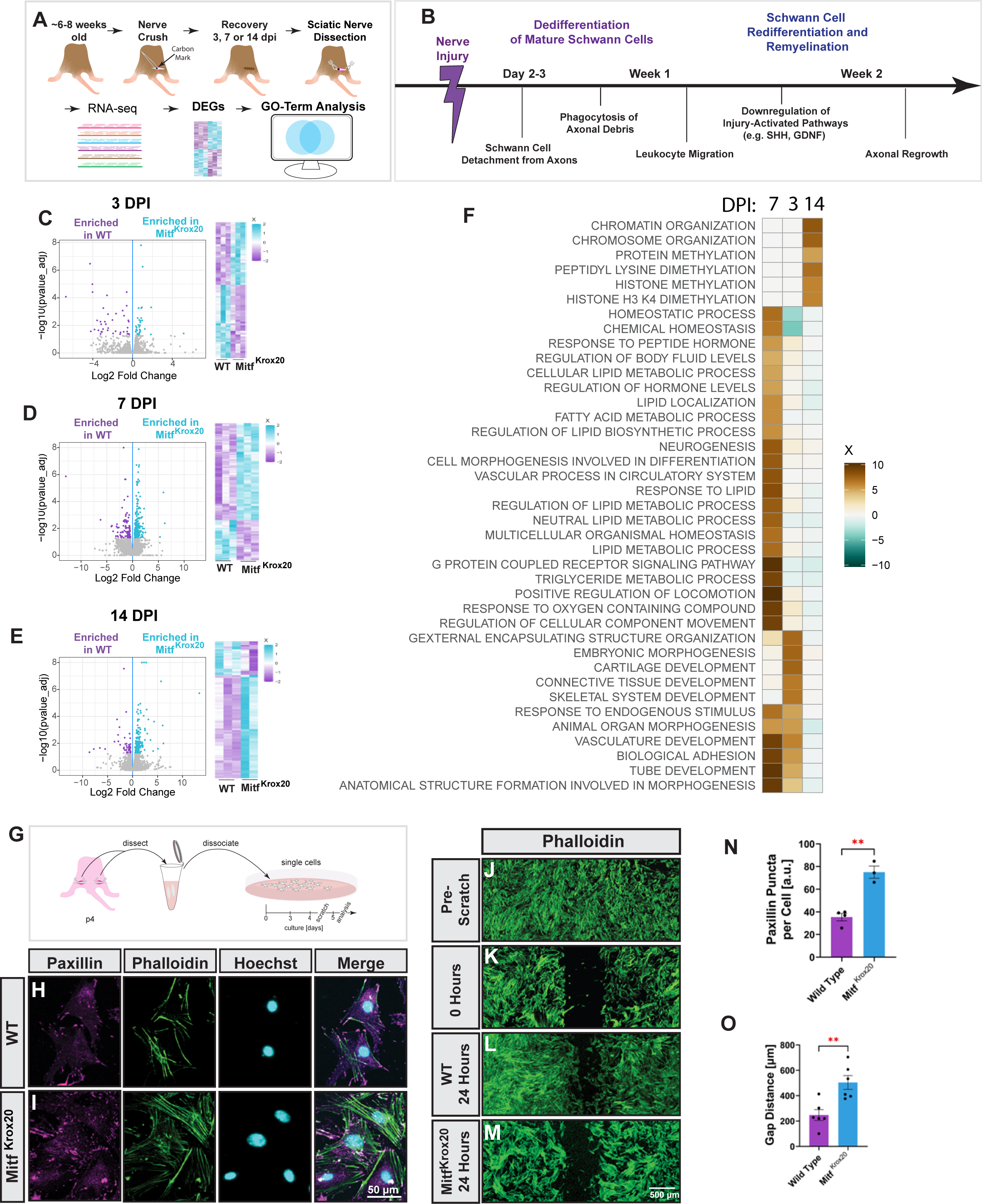
RNA-Sequencing Time Course Reveals A Dynamic Requirement Of Mitf After Peripheral Nerve Injury. **(A)** Workflow for the identification of differentially expressed genes and Gene Ontology in WT and Mitf^Krox20^ animals after injury. **(B)** Time course of major stages of Schwann cell transdifferentiation and redifferentiation after injury. **(C-E)** Volcano plot showing log fold change versus adjusted p-value of sciatic nerve of WT or Mitf^Krox20^ 3 dpi **(C)**, 7 dpi **(D)** or 14 dpi **(E)**. Each dot represents a gene. Blue and purple dots are differentially expressed (n=2-3, each sample was pooled from both sciatic nerves from 3 animals). Heatmap of expression levels (in Z-scores) of differentially expressed genes from. Each row corresponds to a gene. **(F)** Go-term analysis of genes differentially expressed and enriched in Mitf ^Krox20^ 3, 7 and 14 days post injury relative to WT. The top two Biological Process (BP) GO-Term from each cluster is shown. **(G)** Diagram of sciatic nerve dissociation and plating of single cells for Schwann cell primary culture. **(H, I)** Immunofluorescence of primary Schwann cells labeled with Paxillin antibody and stained with Phalloidin-488 and Hoechst from either WT **(E)** or Mitf^Krox20^ **(F)**. **(J-M)** Phalloidin staining of primary Schwann cells grown to confluence **(J)** then scratched **(K)** and allowed to migrate for 24 hours post-scratch, WT **(L)** or Mitf^Krox20^ **(M)** to assess migration. **(N)** Quantification of the number of Paxillin puncta per cell of either WT, or Mitf^Krox20^. At least 40 cells were counted from 3 separate animals (WT) or 4 separate animals (Mitf^Krox20^). Error bars represent S.E.M. Unpaired t-Test. **(O)** Quantification of scratch assay 24 hours after the scratch in either WT or Mitf^Krox20^. Error bars represent S.E.M. Unpaired t-Test.

To determine whether Mitf is required for EMT we designed a functional cellular test to monitor migratory capacity, a key component of mesenchymal behavior (Figure 4G). We found SCs lacking *Mitf* migrated significantly less than WT (Figure 4J-M, O). Downregulation of Paxillin^+^ focal adhesions and the rearrangement of the actin cytoskeleton are critical for the morphological flexibility and migratory capacity of mesenchymal cells. We detected an increase in the number of paxillin^+^ focal adhesions and elevated filamentous actin in *Mitf* mutant SCs (Figure 4H-I, N). Thus, *Mitf* mutant SCs are less migratory and have increased adhesion properties which is counterproductive for EMT and cell migration - key properties of wound healing.

Our sequencing analysis identified transcriptomic changes that impact the dedifferentiation of SCs following injury in *Mitf* mutants. To determine whether these early defects influence SC redifferentiation after injury, we sequenced nerves at 14 dpi, a time point of nascent axonal regrowth and subsequent repair cell redifferentiation into mature SCs. Chromatin rearrangement after injury has been shown to be a powerful factor in generating the repair cell state, in particular H3K4me3 marks sites of injury-related gene expression (i.e. SHH signaling components) ^28, 48^. At 14 dpi, we identified 214 differentially expressed transcripts between WT and Mitf^Krox20^ nerves (188 upregulated, and 26 down-regulated) (Figure 4E). Mitf^Krox20^ mutants had elevated injury-related transcripts including those for histone methylation H3K4me3 and SHH signaling components (Figure 4F, S4C). The continued upregulation of chromatin modifying genes suggests a disruption in the redifferentiation of repair cells back to mature SCs in the *Mitf* mutant.

Taken together, these data reveal that Mitf functions early after injury to regulate a dynamic network of genes that facilitate EMT, metabolism, dedifferentiation of mature SCs into highly specialized repair cells and redifferentiation into mature SCs. We find that these gene expression changes manifest after injury as defects related to repair cell function and plasticity. First, *Mitf* mutants exhibit accelerated axonal degeneration, possibly from a disruption of metabolic support (Figure 3B-F, S3A-D, 4F)^49^. Second, there are early morphological phenotypes of SCs, arising from changes in adhesion properties that disturb membrane flexibility and migratory capacity (Figure 4F-O). Third, these early disruptions to SC dedifferentiation disrupt the subsequent redifferentiation process of repair SCs into mature SCs as evidenced by prolonged injury-related chromatin marks and gene expression (H3K4me3, SHH pathway; Figure 4F, S4C), the persistence of Büngner SCs, excess SC processes in the ECM, ectopic myelination of small caliber axons and wrapping of collagen (Figure 4J-P,S4G0I’). Fourth, these defects in SC transformation following injury diminish axon re-growth and the recovery of sensorimotor function of mice in vivo (Figure 3Q, R).

### mTorc2 Controls Mitf Localization in Schwann Cells

Several kinases have been shown to post-translationally regulate the subcellular localization and transcriptional activity of Mit/Tfe in cell lines (Figure 5A) ^50^. Therefore, we investigated whether Mitf localization was regulated in SCs. To characterize the signaling pathways that control Mitf localization, we established an in vitro primary SC culture system (Figure 5B, Figure S5A-B’’). Antibody staining of endogenous Mitf revealed that it is detected in both the nucleus and the cytoplasm of primary SCs (Figure 5C-C’, O; Figure S5C). To determine whether kinases known to target the Mit/Tfe family alter the localization of Mitf in SCs, we tested antagonists for ERK, PKCβ, GSK3, AKT, mTorc1/2 and mTorc1 ^50^. Inhibitors of ERK1/2 (Sch772984), and its upstream kinase MEK (Trametinib) as well as inhibitors for Akt (MK2206), PKCα/β/γ/ε (Enzstaurin) and GSK3β (BIO) did not alter the subcellular distribution of Mitf (Figure 5A-H’, O). Conversely, treatment of cells with Torin, an mTorc1/2 inhibitor, induced the translocation of Mitf from the cytoplasm into the nucleus (Figure 5I, O). Interestingly, treatment with a selective mTorc1 inhibitor, Rapamycin, did not alter Mitf localization relative to the control (Figure 5N-N’). These data indicate that mTorc2 kinase controls Mitf localization. Interestingly, the Mit/Tfe family member TFEB is activated by mTorc1^44^. Previous studies have shown that mTorc2 kinase activity is stimulated at the plasma membrane through the production of PI(3,4,5)P3 ^51, 52^. To test whether this lipid pathway regulates the localization of Mitf, we inhibited lipid kinases that directly or indirectly contribute to the generation of PI(3,4,5)P3. We inhibited Pikfyve (YM201636) to block the generation of PI(3,5)P2 and Vps34 (VPS34-IN1) to block the generation of PI(3)P. Inhibition of both lipid kinases induced localization of Mitf into the nucleus (Figure 5J-K’, O). These data reveal a lipid kinase signaling cascade that culminates with mTorc2 regulation of Mitf localization (Figure 5P).

**Figure 5.**
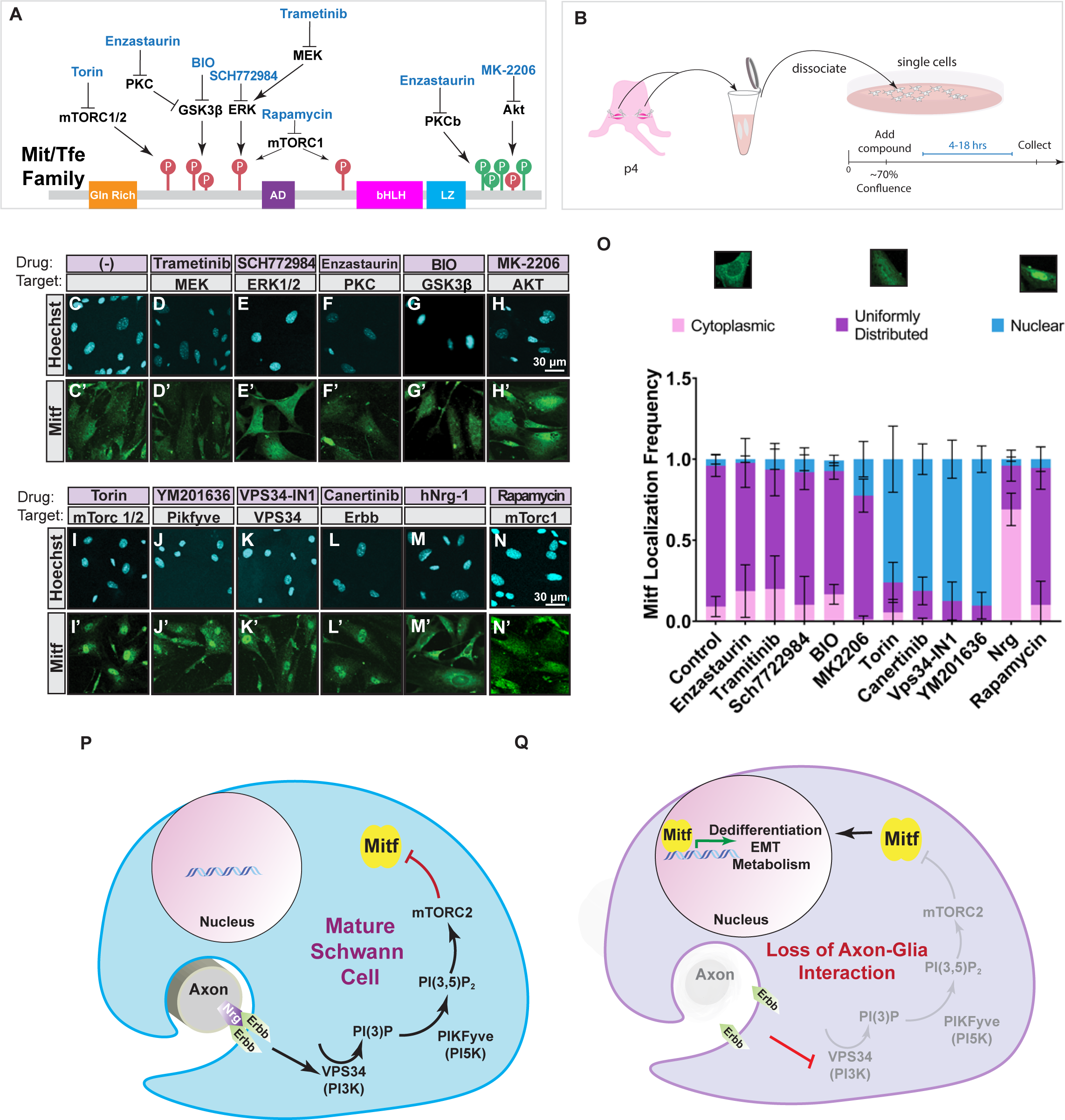
Abrogation of Nrg-Erbb-PI3K-PI5K-mTorc2 Signaling Promotes Mitf Nuclear Translocation. **(A)** Diagram of activating (green) and inhibiting (red) phosphorylation sites (P), of the Mit/Tfe family, the kinases that regulate each of those phosphorylation sites and the pharmacological inhibitors that were used in primary culture to target each of these kinases. **(B)** Diagram of Schwann cell primary culture. **(C-N’)** Primary Schwann cell culture derived from dissociated sciatic nerve of P4 WT (ICR) mice was labeled for Mitf and Hoechst. Primary Schwann cells were cultured with DMSO (Control) **(C-C’)**, Trametinib (MEK) **(D-D’)**, SCH772984 (ERK1/2) **(E-E’)**, Enzastaurin (PKC) **(F-F’)**, BIO (GSK3) **(G-G’)**, MK-2208 (AKT) **(H-H’)**, Torin (mTorc1/2) (**I-I’**), YM201636 (Pikfyve) (**J-J’**), VPS34-IN1 (Vps34) (**K-K’**), Canertinib (ErbB) (**L-L’**), hNrg1 (**M-M’**), Rapamycin (mTorc1) **(N-N’)**. **(O)** Quantification of Mitf enrichment in the cytoplasm (pink), uniformly distributed (purple) or enriched in the nucleus (blue). N= 3 independent experiments for each inhibitor, n>100 cells per experiment. **(P-Q)** Model for Mitf cytoplasmic retention **(P)** and translocation into the nucleus **(Q)**. Abrogation of the Nrg-Erbb-PI3K-PI5K-mTOR signaling pathway results in Mitf protein transport from the cytosol to the nucleus where it activates a gene program that modulates SC dedifferentiation, metabolism, and EMT.

### Nrg is an Axon-to-SC Signal Regulating Mitf

ErbB receptors are expressed by SCs and activate PI3K signaling, therefore we tested whether Nrg1 influenced the localization of Mitf. Addition of ErbB ligand hNrg-1 increased Mitf cytoplasmic retention (Figure 5M-M’, O). Thus, Nrg1 from healthy axons likely activates SC-ErbB receptors and PI3K signaling that suppresses Mitf function by preventing nuclear translocation (Figure 5P). Our findings predict that axonal damage disrupts Nrg1-ErbB signaling thereby increasing nuclear Mitf ^6, 53, 54^. To test this model, we used pan-ErbB inhibitor Canertinib and found it induced nuclear enrichment of Mitf similar to inhibition of PI3K and mTOR (Figure 5L-L’,O, S5C). These data indicate that Nrg-ErbB signaling plays a key role in regulating the subcellular distribution of Mitf in SCs (Figure 5P, Q).

### Peripheral Neuropathy Activates Mitf

In many forms of CMT the physical interaction between Schwann cells and axons are disrupted, raising the question as to whether repair cells are formed and influence disease progression. To investigate Mitf and repair cell function, we created a “tool-box” of conditional Charcot-Marie-Tooth mouse models by targeting genes linked to diseases that cause myelination and axonal degeneration defects. Gene targeting was used to make floxed alleles of: Mtmr2^fl^ (CMT4B), Sh3tc2^fl^ (CMT4C), and Ndrg1^fl^ (CMT4D) (Figure 6A). CMT4B/*Mtmr2* is a dysmyelinating peripheral neuropathy characterized by myelin outfoldings, without major signs of axonal loss ^23, 55^ (Figure 6B-C). CMT4C/*Sh3tc2* is a hypomyelinating disorder with mild, progressive axonal loss ^56^; and CMT4D/*Ndrg1* exhibits hypomyelination that leads to severe axonal atrophy^57^ (Figure 6B,D-E). We found that selective deletion of these genes from SCs in mice using DHH-Cre closely mimicked the human disease pathology of each respective disorder. Namely, Mtmr2^DHH^ displayed hypermyelination of large and small caliber axons, Sh3tc2^DHH^ displayed hypomyelination, and Ndrg1^DHH^ had hypomyelination and axonal loss. Importantly, these conditional gene mutations demonstrate each factor has SC autonomous functions that are required for healthy nerves.

**Figure 6.**
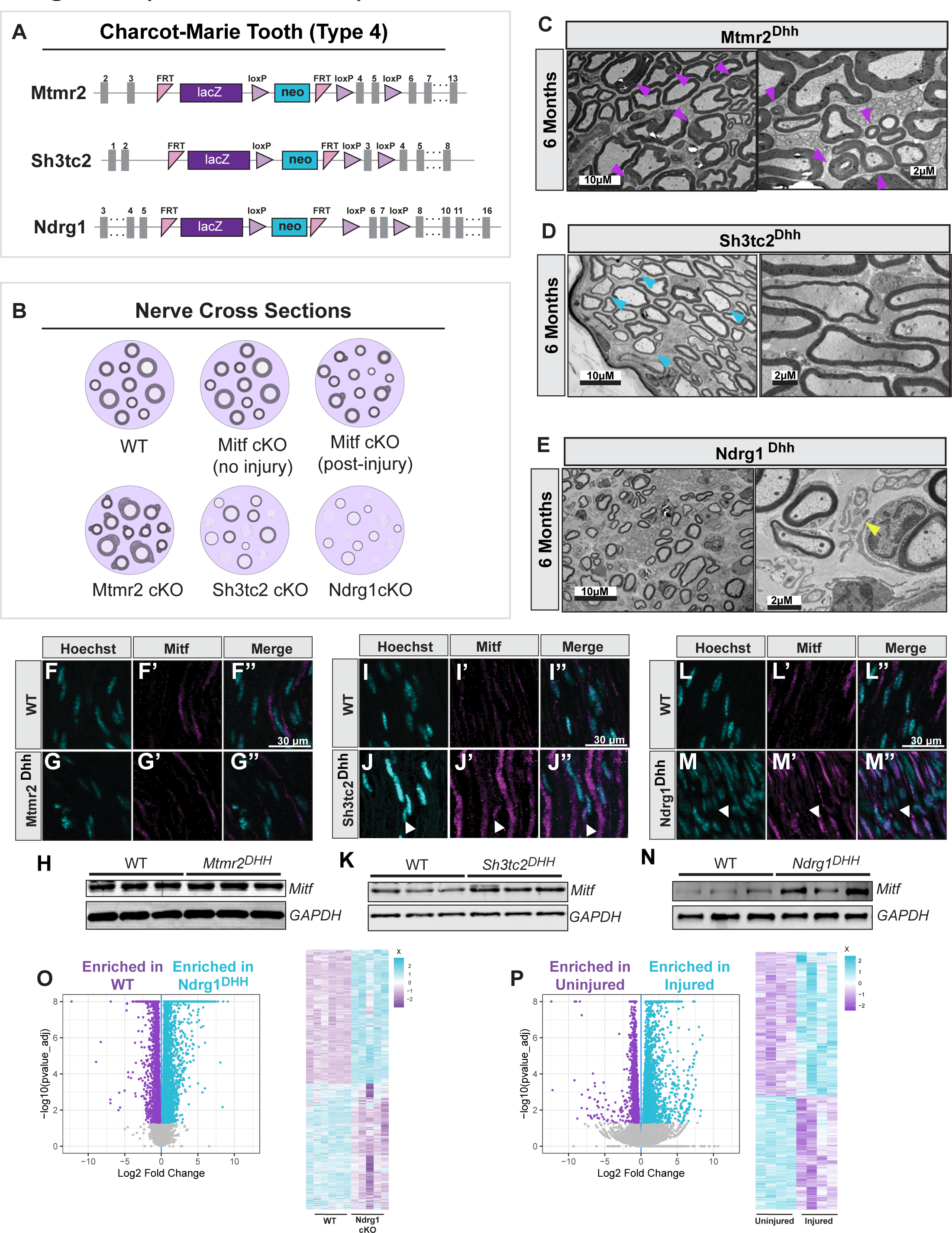
Mitf Is Upregulated In Models Of CMT With Axonal Loss. **(A)** Mtmr2, Sh3tc2 and Ndrg1 knock-in constructs used to generate ‘floxable’ alleles and mouse models of CMT4B, CMT4C, and CMT4D, respectively. **(B)** Diagram of nerve cross sections in WT animals, Mitf^cKO^ before injury, Mitf^cKO^ after injury, CMT4B/Mtmr2^cKO^, CMT4C/Sh3tc2^cKO^, CMT4D/Ndrg1^cKO^. **(C-E)** TEM of Mtmr2^Dhh^ shows myelin outfoldings (**C**, purple arrowheads). TEM of Sh3tc2^Dhh^ and thin myelin (**D,** blue arrowheads). TEM of Ndrg1^Dhh^ reveals the presence of Büngner cells (**E,** yellow arrowheads). **(F-G’’)** Representative immunofluorescence of 6 months old WT (Mtmr2^fl/fl^) or Mtmr2^Dhh^ (Dhh:Cre, Mtmr2^fl/fl^) longitudinal sections of the sciatic nerve double labelled with Mitf and Hoechst. **(H)** Sciatic nerve protein lysates of WT (Mtmr2^fl/fl^) and Mtmr2^Dhh^ analyzed by SDS-page and immunoblotted for Mitf and GAPDH. **(I-J’’)** Immunofluorescence of 6 months old WT (Sh3tc2^fl/fl^) or Sch3tc2^Dhh^ (Dhh:Cre, Sh3tc2^fl/fl^) longitudinal sections of the sciatic nerve double labelled with Mitf and Hoechst. Co- labeling indicated with white arrowhead. **(K)** Sciatic nerve protein lysates of WT (Sh3tc2^fl/fl^) and Sh3tc2^Dhh^ analyzed by SDS-page and immunoblotted for Mitf and GAPDH. **(L-M’’)** Immunofluorescence of 4 months old WT (Ndrg1^fl/fl^) or Ndrg1^Dhh^ (Dhh:Cre, Ndrg1^fl/fl^) longitudinal sections of the sciatic nerve double labelled with Mitf and Hoechst. Co- labeling indicated with white arrowhead. **(N)** Sciatic nerve protein lysates of WT (Ndrg1^fl/fl^) and Ndrg1^Dhh^ analyzed by SDS-page and immunoblotted for Mitf and GAPDH. **(O-P)** Volcano plot showing log fold change versus adjusted p-value of sciatic nerve of Ndrg1^Dhh^ at 4 weeks old compared to WT **(O)** and uninjured compared to injured (3dpi) at 8 weeks old **(P)**. Each dot represents a gene. Blue and purple dots are differentially expressed. Each sample is composed of 6 sciatic nerves from 3 animals. Heat map of expression levels (Z-score) of differentially expressed genes. Each row corresponds to a gene.

Although Mtmr2^DHH^ animals exhibited significant hindlimb and forelimb weakness (Figure S6A-B), Mitf levels and subcellular localization in SCs was normal (Figure 6F-H, S6E). Sh3tc2^DHH^ animals undergo mild axonal loss and display significant hindlimb and forelimb grip strength weakness (Figure S6C-D); Mitf is modestly upregulated in Sh3tc2^DHH^ animals and distributed throughout the cytoplasm and nucleus of SCs (1.3 fold increase, p< 0.05; Figure 6I-K, S6F). Ndrg1^DHH^ animals exhibit severe axonal loss and are significantly weaker than littermate controls (Figure S7G). Mitf protein is elevated in Ndrg1^DHH^ animals and was detected in many SC nuclei compared to controls (4.6 fold increase, p< 0.05; Figure 6L-N, S6G). We find that Mitf is not a readout for the degree of sensorimotor dysfunction, but it does correlate with the breakdown of axon-Schwann cell interaction in the context of disease. The most extreme form of axonal loss is observed in Ndrg1^DHH^ (CMT4D), correlating with the most dramatic increase in the overall levels and nuclear localization of Mitf.

### Shared Transcriptomic Response in Injury and Chronic Disease

The increase in Mitf protein concentration and nuclear localization in Ndrg1^DHH^ (CMT4D) is reminiscent of Mitf dynamics following nerve injury (Compare 6N to Figure 2K). We therefore compared the transcriptomic changes of the nerve from Ndrg1^DHH^ mice to transected nerves from WT mice. Intriguingly, both injured and Ndrg1^DHH^ nerves downregulated factors linked to cellular metabolism, cytoskeleton, cell junction organization and adhesion (Figure 6O-P, S6H-J). Similar results were observed when analyzing the pathways linked to downregulated gene expression of injured and Fig4^plt^ (CMT4J) nerves (Figure S7A). Similar to observations in injury, the transcriptomic profiles of CMT4D and CMT4J are consistent with an increase in mesenchymal characteristics and a switch from mature SCs to dedifferentiated repair cells.

Given the remarkable degree of shared differential gene expression between these pairwise comparisons, we examined whether there was a core transcriptomic program that was shared across injury and neural disease. We found a significant degree of shared differentially expressed genes between injury (3dpi), CMT4D (Ndrg1^Dhh^) and CMT4J (Fig4^plt^) (571 genes, Figure 7A-B). Several themes emerged from the GO-Term analysis of these shared genes. This included: (1) Regulation of cell migration, adhesion and cell morphogenesis, hallmarks of the required epithelial to mesenchymal transition that mature Schwann cells undergo to form functional repair cells. (2) Changes in glial cell differentiation and nervous system development consistent with a SC transdifferentiation into a repair cell. (3) Changes to the lipid, sterol and alcohol biosynthetic process consistent with changes in cellular metabolism as mature SCs transform into repair cells (Figure 7C). These data suggest the existence of a “core” 571 gene transcriptional response of the peripheral nerve that is shared between injury and chronic disease (CMT4D and CMT4J).

**Figure 7.**
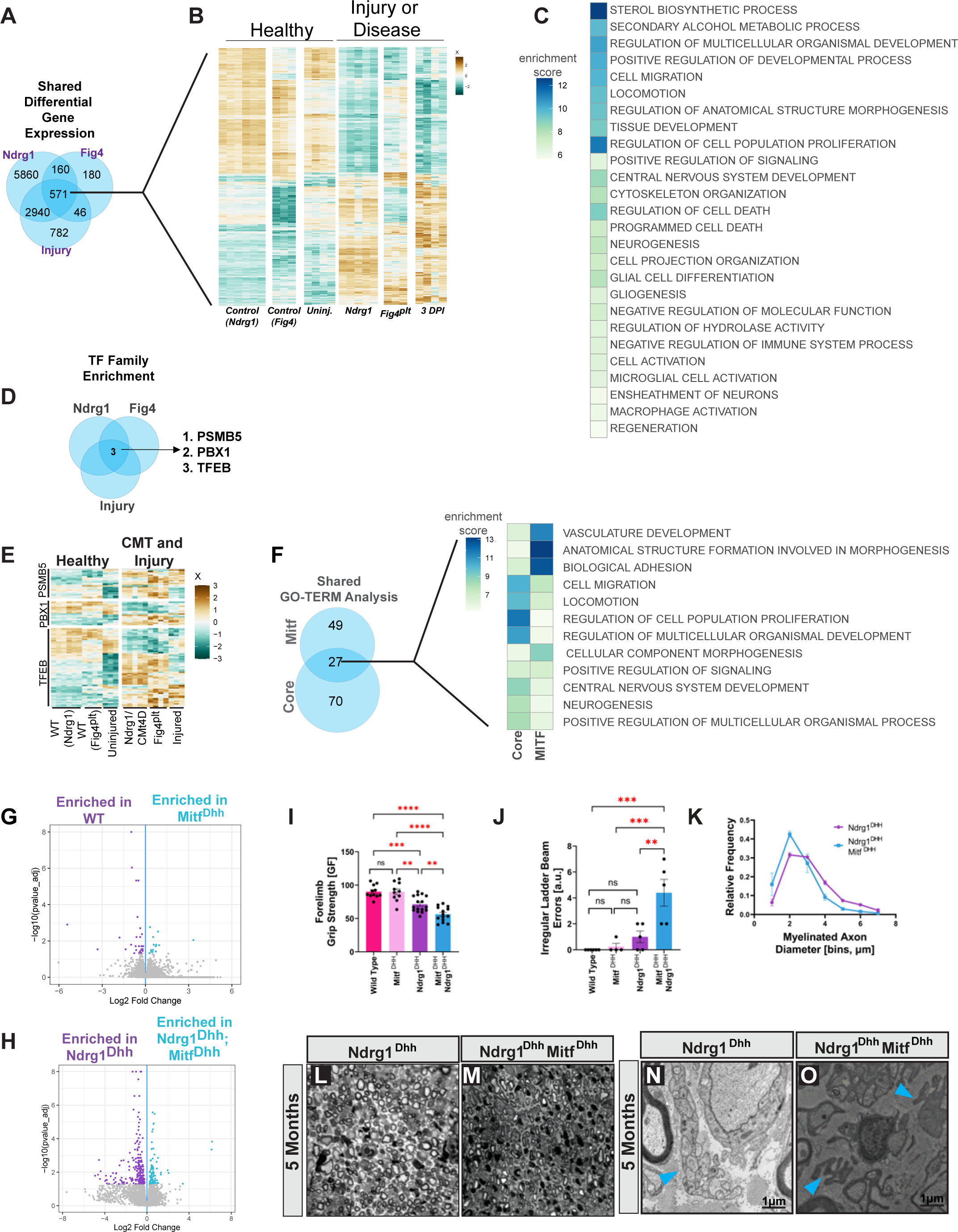
Mitf Facilitates Core Nerve Response Pathways to Attenuate Disease. **(A)** Venn Diagram depicting the shared differential gene expression between Ndrg1^Dhh^, Fig4^plt^ and injured nerves (3dpi). 571 genes are differentially expressed in all three categories. **(B)** Heat map of expression levels (Z-score) of the 571 differentially expressed genes shows levels of differential expression between genes compared to their respective control (Ndrg1^Dhh^, Fig4^plt^ and injured nerves (3dpi)). **(C)** GO-Term analysis of the shared 571 genes differentially expressed in Ndrg1^Dhh^, Fig4^plt^ and injured nerves. The top two Biological Process (BP) GO-Term from each cluster is shown. **(D)** GSEA prediction of transcription factors enriched in the 571 shared differentially expressed genes from **(A)**. **(E)** Heat map of expression levels (Z-score) of differentially expressed genes predicted to be regulated by PSMB5, PBX1 or TFEB transcription factor families. **(F)** Venn Diagram of GO-Terms from differential expression of Mitf^Krox20^ (3 and 7 dpi) compared to the GO-Terms from the shared gene expression, termed ‘Core’ (from **C)**. Heat map showing the top two Biological Process (BP) GO-Terms from each cluster is shown. **(G)** Volcano plot showing log fold change versus adjusted p-value of sciatic nerve of Mitf^Dhh^ at 4 weeks old compared with littermate controls. Blue and purple dots are differentially expressed genes (N= 5, each sample pooled from both sciatic nerves of 3 animals). **(H)** Volcano plot showing log fold change versus adjusted p-value of sciatic nerve of Ndrg1 cKO at 4 weeks old compared with Ndrg1^Dhh^; Mitf^Dhh^. Blue and purple dots are differentially expressed genes (N= 5, each sample pooled from both sciatic nerves of 3 animals). **(I)** Forelimb grip strength of control (n=11), Mitf^Dhh^ (n=10), Ndrg1^Dhh^ (n=16) and the double conditional knockout Ndrg1^Dhh^Mitf ^Dhh^ (n=14) animals at 8 weeks. Ordinary One-way ANOVA with Tukey’s multiple comparison test. **(J)** Irregular ladder Beam Errors of WT (N=5) Mitf^Dhh^ (N=4) Ndrg1^Dhh^ (n=5) Ndgr1^Dhh^; Mitf^Dhh^ (N=5). Ordinary One-way ANOVA with Šidák’s multiple comparison test. **(K)** Quantification of the frequency of myelinated axon diameter in Ndrg1^Dhh^ and to Ndgr1^Dhh^; Mitf^Dhh^ sciatic nerve at 5 months old (n=3, >150 axons per animal). Multiple unpaired t-Test. **(L, M)** Representative Toluidine Blue stained cross sections of sciatic nerve from Ndrg1^Dhh^ (C) or Mitf^Dhh^Ndgr1^Dhh^ littermates. **(N, O)** Representative TEM images of Büngner Band formation in 5 months old Ndrg1^Dhh^ (N) and Ndgr1^Dhh^Mitf^Dhh^ (O) littermates in the sciatic nerve (blue arrowheads). (Büngner Bands were not present in the control or Mitf^Dhh^ single mutant).

### Mitf Regulation of Core Programs Activated by Nerve Injury and Disease

To identify potential regulators of the 571 “core” genes in injury and disease, we identified transcription factor binding motifs enriched in their promoter regions in the injury, CMT4D (Ndrg1^Dhh^) and CMT4J (Fig4^plt^) mice. GSEA revealed motif enrichment of three transcription factor families represented by: PSMB5, PBX1 and TFEB (Mit/Tfe family which includes Mitf) (Figure 7D- E). Next we compared the “core” pathways predicted to be regulated by Mit/Tfe family members to the pathways misregulated in the *Mitf* mutants following injury. We found that the pathways Mitf regulates in SCs after injury overlap with those key pathways from the “core” transcriptomic response (Figure 7F). Pathway overlap between these two groups include: cell migration, adhesion and cellular morphogenesis, metabolic processes and differentiation. Taken together, these findings reveal a Mitf regulated gene set activated during trauma that functionally overlaps with disease thereby defining a core gene set associated with axonal degeneration.

### Mitf Attenuates Disease in vivo

Mitf is predicted to regulate a core program of changes linked to nerve repair during chronic disease. Therefore, we sought to determine the role that Mitf plays in the manifestations of CMT4D disease. We performed RNAseq on sciatic nerves from WT, Mitf^DHH^, Ndrg1^DHH^, and Ndrg1^DHH^ :Mitf^DHH^ mice. The transcriptomes of WT compared to uninjured Mitf^DHH^ nerves revealed few differences (14 upregulated, 29 downregulated genes relative to the control; Figure 7G, S7B). In contrast, comparison of the sciatic nerve transcriptome from Ndrg1^DHH^ animals to Ndrg1^DHH^ Mitf^DHH^ identified 222 upregulated and 85 downregulated genes (Figure 7H, S7C). GO-Term analysis of differentially expressed genes revealed dysregulation of pathways linked to adhesion, cell migration and homeostasis in Ndrg1^DHH^ :Mitf^DHH^ nerves (Figure S7D-F).

While Mitf^DHH^ animals do not exhibit grip strength phenotypes, we discovered that Ndrg1^DHH^ :Mitf^DHH^ were significantly weaker than single Ndrg1^DHH^ mutants (Figure 7I, S7G). We next assayed motor coordination using an irregular ladder beam. Uninjured WT and Mitf^DHH^ animals were coordinated and rarely missed rungs, whereas Ndrg1^DHH^ mice had detectable missteps. The Ndrg1^DHH^ :Mitf^DHH^ animals performed significantly worse than the Ndrg1^DHH^ mutants (Figure 7J, S7H-K). To identify potential cellular defects in these mutants we analyzed TEM of the sciatic nerve. The organization of SCs and axon size distribution was similar between wild type controls and (uninjured) Mitf^DHH^ mutants. We found that Ndrg1^DHH^ :Mitf^DHH^ animals had fewer large caliber axons compared to Ndrg1^DHH^ mutants (Figure 7K-M). As expected, Büngner SCs were absent from control and Mitf^DHH^ sciatic nerves; however, they were frequently observed in Ndrg1^DHH^ and Ndrg1^DHH^ :Mitf^DHH^ animals, consistent with axonal loss (Figure 7N,O). While Büngner SCs in Ndrg1^DHH^ were round, the Ndrg1^DHH^ :Mitf^DHH^ animals exhibited a flat morphology, consistent with a loss of regenerative capacity of the Büngner cells ^12, 58^. Given that (uninjured) Mitf^Dhh^ animals were indistinguishable from WT, the behavioral and cellular phenotypes arising from Ndrg1^DHH^ :Mitf ^DHH^ double mutants are not simply additive of those in the single mutants. These data reveal that Mitf has a compensatory function that attenuates the natural progression of CMT4D in vivo and is a key regulator of gene expression induced by this disease.

## Discussion

Physical damage to peripheral nerves is efficiently repaired by SCs that detect axonal damage and transiently transform into repair cells that reestablish myelinated neuronal connections - thereby restoring sensorimotor function. In this study we identify the transcription factor Mitf as a critical regulator of SC transdifferentiation and repair cell function whose subcellular localization is coupled to the Nrg1 signaling pathway. An important discovery that emerged from the molecular characterization of SC plasticity was uncovering the role Mitf and repair cells play in limiting the severity of inherited peripheral neuropathies such as CMT4D. We discuss how SCs detect axonal damage leading to activation of a core genetic program controlled by Mitf that counteracts nerve damage from injuries and diseases.

### Mitf is a sensor of axonal damage

Although Mitf is constitutively expressed by mature SCs, its transcriptional activity appears to be linked to post-translational modifications that regulate its subcellular localization. Our findings suggest mature SCs are readily poised to switch their gene expression in response to nerve damage by activating a cytoplasmic-to-nuclear redistribution of Mitf. To understand how Mitf localization is gated we probed a variety of signaling pathways using pharmacological agents. We found that the Nrg1 ligand, expressed by sensory and motor neurons, activates ErbB receptors on SCs and this inhibits Mitf. Thus the interaction between SCs and healthy axons triggers Nrg1- ErbB signaling that holds Mitf in a transcriptional “off” state within the cytoplasm of SCs (Figure 5P). Consistent with previous characterization of Nrg1-ErbB downstream signaling, the cytoplasmic retention of Mitf is dependent upon a series of lipid second messengers produced by PI3K and PI5K, which we found control the mTORC2 variant of the mTOR serine/threonine kinase complex in SCs (Figure 5P). Consequently axon damage is predicted to reduce Nrg1-ErbB signaling, leading to downregulation of mTORC2 activity and nuclear translocation of Mitf (Figure 5Q). Interestingly, Mitf protein levels increase following nerve injury, whereas mRNA levels remain constant. Thus, Nrg1 signaling likely controls both the localization and the half-life of Mitf protein.

Mitf is a member of the Mit/Tfe family of transcription factors, which includes Tfeb, Tfec, Tfe3, and Mitf, expressed in varying combinations across many cell types ^38^. For example, Tfeb is enriched in oligodendrocytes, the myelinating glia of the central nervous system that serve as the counterpart to SCs in the peripheral nervous system ^39^. While it is tempting to compare Tfeb and Mitf functions, Tfeb regulates oligodendrocyte cell number during development^39^, whereas we found Mitf is dispensable for SC development but is required for nerve repair. Interestingly Mit/Tfe transcription factors have been linked to the regulation of lysosomal function and autophagy ^44, 46, 50, 59^, but neither Tfeb in oligodendrocytes ^39^ nor Mitf in SCs was found to be required for these functions. In the context of SCs this apparent discrepancy could be due to transcriptional redundancy because Tfeb, Mitf and other family members are co-expressed and may therefore cooperatively regulate autophagy. The disruption of autophagy following nerve injury in calcineurin mutants indicates this phosphatase may contribute to the activation of multiple Mit/Tfe family members in SCs ^60^.

The *Nrg1* gene has many splice variants that encode functionally distinct proteins across a variety of cell types ^61^. Type I Nrg1 is upregulated within SCs after injury ^62^, raising the question of how this factor could be a signal of axon damage that controls Mitf localization? Potentially the concentration, splice variant form, and/or autocrine versus paracrine signaling influence how Nrg1 functions during nerve injury. In addition, there may be a temporal order in which nerve injury transiently disrupts Type III Nrg1-ErbB signaling causing Mitf to trigger the formation of repair SCs, followed later by an upregulation of Type I Nrg1 within repair cells to promote the reformation of myelinating SCs ^62–64^. Although there are additional details of Nrg1 signaling to define, recent studies have shown that the inappropriate upregulation of Nrg1 in SCs causes hypermyelination ^65^. Our studies indicate Mitf activity will be dysregulated when Nrg1 is either under- or over- expressed, impacting how SCs function and respond to environmental cues.

### Mitf’s role in Schwann cell plasticity

Peripheral nerve recovery from injury is dependent upon the ability of SCs to transform into repair SCs ^1^. The functions of repair cells include metabolic support of axons, macrophage recruitment, clearance of cellular debris, and formation of Büngner scaffolds ^5^. After axons regrow, repair cells revert back into mature SCs that remyelinate axons and thereby restore efficient nerve conduction necessary for sensory and motor function ^6^. Interestingly, some of these repair steps can be dissociated with genetics. The mutation of *Mitf* causes an accelerated degradation of injured axons and aberrant remyelination of new axons - but myelin debris is removed normally. In contrast, SCs lacking *c-Jun* fail to clear myelin debris following nerve injury although Büngner cell formation and axonal regrowth are preserved ^12^. These genetically-separable functions support the view that the transformation of SCs into repair cells followed by the retransformation of repair cells back into myelinating SCs is a tightly-choreographed sequential process. However, it also remains possible that mature myelinating and non-myelinating Remak SCs, for example, transform into separate subpopulations of repair cells following injury that have distinct functions specified by different genetic pathways.

In the absence of Mitf, we found damaged axons degenerate more rapidly, possibly because the glycolytic activity of Mitf-deficient repair SCs is compromised ^49, 66^. In addition, *Mitf* mutants displayed striking alterations in remyelination following injury. It is not known why remyelination is defective, however we found that chromatin organization genes became dysregulated in *Mitf* mutants and this might impair the gene expression necessary for conversion of repair cells back to myelinating SCs. Taken together, our studies indicate that Mitf is necessary for two critical aspects of SC plasticity: the transformation of mature SCs into functional repair cells that support axonal integrity, and the redifferentiation of repair cells into functional myelinating SCs that restore normal nerve conduction.

### Nerve homeostasis and core genetic programs

Peripheral neuropathies arise from gene mutations in *Fig4* (CMT4J), *Ndrg1* (CMT4D), *SH3TC2* (CMT4C), and *MTMR2* (CMT4B). These genes directly encode and/or regulate phosphatases, kinases, and GTPases involved in cellular signaling and membrane trafficking. We found that selectively deleting the *Ndrg1* and *SH3TC2* genes in SCs led to sensorimotor defects (i.e. peripheral neuropathies), indicating that each of these factors functions intrinsically within SCs and is required for maintaining healthy nerves. When we compared the transcriptional changes in nerves from *Fig4* and *Ndrg1* cKO mutants to the gene expression in injured nerves we discovered a core set of 571 genes shared across each condition. The predicted functions of this core set of common genes were cell de-differentiation, metabolism, and migration. Thus, the gene expression alterations arising from chronic degenerative conditions such as CMT4J (*Fig4* cKO) and CMT4D (*Ndrg1* cKO) share striking similarities to the genes activated by acute nerve injury. The promoters of this core gene set where enriched for Mit/Tfe binding sites, suggesting Mitf may contribute to their regulation. This was further supported by the finding that mutation of *Mitf* caused sensorimotor deficits to become more severe in CMT4D (*Ndrg1* cKO).

Because Mitf is necessary for the generation of repair cells and the deletion of Mitf makes neurodegeneration worse in CMT4D (*Ndrg1* cKO), it is likely that repair cells are actively recruited and provide a functional benefit during the course of this disease. Moreover, these findings raise the possibility that SC transdifferentiation is a process that is also needed to maintain healthy nerves by replacing support cells and repairing myelin as needed. Consequently the signaling defects caused by genetic mutations in CMT4B/C/D/J may alter Mitf-activation, thereby impacting repair cell generation and nerve homeostasis. Thus, the core gene program controlled by Mitf including factors involved in de-differentiation, metabolism, and migration may represent targets that counteract the effects of aging, injury and disease on peripheral nerve function.

## Materials and Methods

### Mouse lines

Plasmids containing the Mitf knock-in allele were obtained from the Knockout Mouse Project (KOMP) with Mitf exon 6 (a common exon shared between splicing variants) flanked by LoxP sites (project number: 29766; Plasmid ID: PRPGS00031_A_F01). Mitf conditional animals (Mitf^FL^) were generated by homologous recombination using the “knockout first” tm1a plasmid available at komp.org which inserts a construct to create either a LacZ reporter or a FLP-dependent conditional knock out. Embryonic stem cells were electroporated with the linearized Mitf^FL^ targeting vector, and colonies were screened by PCR for insertion and confirmed by Southern blot using an external 5’ probe. Two properly targeted clones were injected into blastocysts, followed by implantation. Upon birth, the chimeric progeny was analyzed for germline transmission, and gene insertion expression was confirmed by detection of LacZ in immunoblots. Progeny were crossed to Actb:Flp (from M. Goulding, Salk), to excise the LacZ/Neo selection cassette, resulting in LoxP sites flanking exons 4 and 5 of Mitf, generating the Mitf^FL^ allele.

Null Mitf (Mitf cKO) were generated in Mitf^FL/FL^ mice mated to either Dhh:Cre (*JAX Cat# 012929)* or Egr2:Cre (Jax *Cat# 025744) mice* and maintained on a mixed C57BL/6; ICR background. Ndrg1 conditional animals (Ndrg1^FL^) were similarly generated by homologous recombination using the “knockout first” tm1a plasmid available from komp.org (plasmid ID# PG00097_Z_4_F03). Crossing Ndrg1^FL^ to Actb:Flp results in LoxP sites flanking either side of exon 2 of the Ndrg1 gene locus. A similar strategy was used to generate Sh3tc2^fl^ alleles (Sh3tc2 KOMP plasmid ID PRPGS00085_A_B05). Germplasm was obtained from KOMP to generate Mtmr2^fl^ mice. Genotyping for the floxed allele is performed by Transnetyx (Transnetyx.com). *Krox- 20 Cre* was obtained from Jax *(Cat# 025744). Ai75 (RCL-nlsT)-D* was obtained from Jax *(Cat# 025106).* ICR obtained from Jax. Mice were housed up to 5 per cage, kept in a 12 h light-dark cycle, and fed standard chow *ad libitum*. Mice were euthanized using CO2. All animal procedures were approved by the Salk Institute for Biological Studies Veterinary Staff, and conducted according to our IACUC protocol.

### Antibodies, Agonists and Chemical Inhibitors

hNrg1-1 (25 ng/ml) (Cell Signal Technology, Catalog # 5218SC); VPS34-IN1 (1 µM) (EMD Millipore, Catalog # 5326280001); YM201636 (2 µM) (Cayman Chemical, Catalog # 13576-1); Canertinib (5 µM) (Fisher Scientific, Catalog # 508629); MK-2206 (1 µM) (Fisher Scientific, Catalog # NC0709830); Torin (5 µM) (Fisher Scientific, Catalog # 501014432); Enzstaurin (5 µM) (Fisher Scientific, Catalog # 501360879); Trametinib (5 µM) (Fisher Scientific, Catalog # NC0991754); KT5720 (VWR International, Catalog # 80512-534); Sch772984 (5 µM) (Fisher Scientific, Catalog # 501152277); BIO (10 µM) (GSK3, Sigma-Aldrich Inc Cat# B1686); Rapamycin (0.27 µM) (mTorc1,Fisher Scientific, Catalog # AAJ67452XF). Rabbit polyclonal Mitf antibody was raised in house (Animal # 7414, # 7416) against antigens DLVNRIIKQEPVLENCSQE (N-term) and CGTMPESSPAYSIPRKMGSNLEDILMD (C-term), respectively, each independently conjugated to KLH (Fisher Scientific, Cat# PI77671). Commercial antibodies: anti-Paxillin (BD Biosciences #610051). Anti-Mitf (Abcam, ab122982), anti-GAPDH (Fitzgerald Cat # 10R-G109a), anti-GFAP (Aviva Cat # OAPC00115, anti-GFAP (Aviva Cat# OAEB01041). Alexa-conjugated secondary antibodies (Life Technologies).

### Western blots

Sciatic nerves were quickly dissected after euthanasia and placed on dry ice prior to lysis or preservation at -80 °C. Protein lysates were generated by nerve homogenization in 50mM HEPES, pH 7.4, 300mM NaCl, 1% Tx-100 with 0.5% Protease Inhibitor Cocktail (Sigma # P8340). Lysates were centrifuged at 4°C at 15,000 RPM for 15 minutes in a table top centrifuge (Eppendorf 5424R). The supernatant’s protein concentration was measured using Bradford’s assay (Bio-Rad #5000202), and adjusted to 1 µg/µL with lysis buffer. Samples were denatured in Laemmli Sample Buffer (5X : 10% SDS, 50% Glycerol, 312mM Tris-HCL pH 6.8, 2% β- Mercaptoethanol) and heated to 100°C for 5 minutes. 5-10 µg of total protein was analyzed by SDS-polyacrylamide gel electrophoresis (SDS-PAGE) followed by immunoblot with in-house rabbit anti-Mitf antibody (1:250) and mouse anti-GAPDH (1:5000) followed by DyLight or Alexa-conjugated secondary antibodies (1:10000) and scanning on an Odyssey CLX scanner (Li-Cor Biosciences). Image quantification was performed on Odyssey software (Li-Cor Biosciences).

### Immunohistochemistry

Sciatic nerves were freshly dissected after euthanasia and fixed by immersion in 4% paraformaldehyde (Electron Microscopy Sciences # 15713) for one hour, followed by three washes in 1X PBS and immersion in 1x PBS + 30% sucrose for 48 hours at 4°C. Sciatic nerves were embedded in OCT (Tissue-Tek #4583), frozen at -20°C and sectioned at 12-20 µm thickness onto glass slides (Fisherbrand 12-550-15) and air dried. Primary Schwann cells were grown on poly-D-Lysine (PDL) coated coverslips (Corning #354085) at 37°C with 5% CO2 and then fixed with 4% paraformaldehyde in 1x PBS for 10 minutes at room temperature followed by three 1x PBS washes. Primary and secondary antibodies were diluted in 1x PBS, 20% Donkey Serum (Sigma #D9663), 0.5% Tx-100 and then sterilized by filtration. Abcam Mitf rabbit polyclonal 1:500. Aviva anti-GFAP Chicken Polyclonal (1:2000). Aviva Anti-GFAP Goat Polyclonal (1:2000). Anti-Paxillin Mouse Monoclonal (1:500). Assorted secondary antibodies from Life Technologies were used to match the host species of the primary antibody at 1:500 dilution. Cellular Stains: Alexa Fluor 488 Phalloidin (Life Technologies #A12379, 1:1000); Fluoromyelin (Life Technologies #F34651, 1:1000), Hoechst 33342 (Life Technologies #H3570, 1:2000). Fluorescence Microscopy: Images were acquired on an Olympus Fluoview 3000 point-scanning confocal microscope with 4 lasers and 2 GaSP detectors and a hybrid resonant-galvanometer scanning mirror and specialized long working distance 20X objective.

### Surgical Procedures

A blend of fentanyl (0.05 mg/kg), Midazolam (5 mg/kg), Medetomidin/Dormitor (0.5 mg/Kg) were used as anesthesia for all procedures. The unilateral or bilateral nerve crush was conducted by crushing the sciatic nerve below the sciatic notch using cooled forceps and marking the crush site with carbon. Nerve transections were cut at the sciatic notch. After transection, the distal stump was positioned 1-2 mm away from the proximal nerve so that axons could regrow through the distal stump. For analgesia, 0.1 mg/kg buprenorphine SR was injected intraperitoneally immediately following the surgery, before the animals were revived with a blend of Flumaxenil/Anexate (0.5 mg/kg) and Antisedan (2.5 mg/Kg). Ibuprofen was administered (0.112 mg/ml) in the water for up to 7 days following surgery. Animals were monitored in accordance with our IACUC protocol.

### Electron Microscopy

Sciatic nerves were freshly dissected after euthanasia and immersed in Karnovsky’s fixative (2.5% gluteraldehyde and 4% paraformaldehyde in 0.1M Cacodylate buffer) and incubated overnight at 4°C. The next day samples were washed 3 times in 0.1M Na Cacodylate, pH7.4 for 15 minutes. Samples were then incubated in 2% osmium tetraoxide, 0.1M Na Cacodylate, for 1 hour at room temperature (RT) in the dark followed by 3 washes 15 min each with molecular biology grade water at RT, and stained with 2% Uranylacetate for 1 hour in the dark at RT. Samples were serially dehydrated from 25% ethanol to 100% ethanol and then submerged in 100% acetone for 15 minutes, followed by immersion in a mixture of 2:1 Acetone:EPON for 1-2 hours at RT. The samples are left overnight in 1:1 Acetone:EPON, followed the next day by incubation in fresh 100% EPON for 4 hours at RT and finally embedded overnight in EPON at 65°C. Fixatives and embedding chemicals are from Electron Microscopy Sciences. EPON resin was mixed according to the manufacturer specifications.

### Morphological Analysis

Semi-thin sections (500 nm) were stained with 1% toluidine blue and used for myelinated axon counts. Ultrathin sections (45-50 nm) were imaged with an FEI Tecnai Spirit G2 BioTWIN Transmission Electron Microscope equipped with an Eagle 16 megapixel camera. Random images were acquired for analysis. To calculate G-Ratio (i.e. the ratio between axon diameter and axon plus myelin diameter), the axon diameter and fiber diameters were measured using FIJI. At least 100 fibers per nerve were analyzed.

### Animal Behavior

*<u>The Hargreaves Assay</u>* was conducted using male and female mice habituated in the testing environment for 2 hours before the day of the assay, and 2 hours on the day of the assay. Animals were then placed on a standard Hargreaves glass platform, where they were set in single isolation. Either the left (uninjured) or the right (injured) hind paw was measured once every 5 minutes. Animals were only measured if they were inactive before the test began. Lamp power was set to 30%. Weekly measurements were conducted before nerve crush, and then each week for 8 weeks after nerve crush. *<u>The Rotorod Assay</u>* was conducted using a standard rotorod program accelerating from 0-30 rotations per minute over five minutes; the latency time for each animal was measured over 4 trials. <u>The Grip Strength Assay</u> was conducted using male and female animals. Both hindlimbs or both forelimbs were tested together using a standard grip strength apparatus. For some animals (Figure 3) either the right (injured) or the left (uninjured) hindlimbs were tested for strength individually. <u>The Ladder Beam Assay</u> was performed by a blind observer who acclimated the mice to the ladder by allowing them to walk across with all prongs inserted. Designated ladder rungs were then removed, and mice were allowed to walk across the ladder 4 times. Simultaneous video capture of both the side and belly profiles were obtained. Slips were counted when an animal’s hindlimb completely missed a ladder rung, such that the entire leg falls below the level of the ladder.

### Primary Schwann Cell Culture

To prepare primary cultures of mouse Schwann cells, sciatic nerves were dissected at P4 from ICR wild type animals and cultured to ∼50% confluence, typically taking 2-4 days. (Higher rates of confluency corresponded to Schwann cell clumping and detachment from the wells). The nerves were placed into warm Dissection Media (Low Glucose-DMEM (Life Technologies), 0.025M HEPES (Life Technologies), Penicillin/Streptomycin (Life Technologies), and antibiotic/antimyotic (Life Technologies)). Nerves were then treated with 1.25 mg/ml trypsin and 2 mg/ml collagenase for 20 minutes at 37°C. Nerves were washed in Feeding Media (10% FBS (Life Technologies), Low Glucose-DMEM (Life Technologies), 0.025M HEPES (Life Technologies), Penicillin/Streptomycin (Life Technologies), and antibiotic/antimyotic (Life Technologies)). Nerves were then triturated through an 18-gauge and then a 21-gauge needle. Epineurium and debris were removed with sterile tweezers and then remaining cells were plated onto PDL-coated coverslips in 50 µL droplets. Cells were allowed to adhere to the coverslip for 1- 2 hours and then additional Feeding Media was added to submerge the cells. <u>Agonist/Antagonist</u> <u>Assay</u>: Typically, cells grew for 3-5 days before the addition of drugs. Agonists and Inhibitors were added at 50% confluency for 18 hours, except for BIO (GSK3 inhibitor) which was incubated for 5 hours. <u>Scratch Assay:</u> Schwann cells were grown to 100% confluency as determined by light microscopy. A pipette was then applied to the coverslip to remove cells from the center of the coverslip. Cells were incubated for 24 hours, washed with 1x PBS, and fixed with 4% paraformaldehyde and stained with 66 µM phalloidin-488 at 1:500 for 1 hour at RT in the dark.

### Nerve Explants

Distal Sciatic Nerve segments from Egr2:Cre, Ai75 (Lox-Stop-Lox- nuclear Td- tomato reporter) mice were quickly dissected and either immediately fixed (time point 0, uninjured condition) or transferred to warm Feeding Media (Low glucose DMEM with 10% Fetal bovine serum, 0.025M HEPES, Penicillin/Streptomycin, and antibiotic, antimyotic) and cultured in 5% CO2 at 37°C. After incubation, nerves were gently washed once with 1x PBS and fixed in 4% paraformaldehyde for 1 hour at RT and then washed in 1x PBS.

### RNA-sequencing

High-throughput whole transcriptome mRNA-Sequencing: Sciatic nerves were rapidly dissected after euthanasia, and quick-frozen on dry ice. Sciatic nerves were lysed with a pellet pestle in RLT buffer (Qiagen) with β-mercaptoethanol. Total RNA was isolated using Qiagen RNAeasy Plus Micro kit (# 74034). The Agilent Bioanalyzer (Tape Station) was used to determine RNA integrity (RIN) prior to library preparation. Stranded mRNA sequencing libraries were prepared using the TruSeq Stranded mRNA Library Prep Kit according to manufacturer’s instructions (Illumina). Libraries were then quantified, pooled, and sequenced at paired end 75 (PE75) base-pair using the Illumina NexSeq (150 cycle, high output) at the Salk Institute for Biological Studies Next Generation Sequencing Core (Figure 1, 4, S1, S4; Fig4 developmental and Mitf injury data) or the University of California, San Diego Genomic Sequencing Core (Figure 7, S7 Mitf and Ndrg1 data). <u>Analysis:</u> Single and paired-end RNA-Seq reads were quantified using Kallisto against the Gencode (M17) transcript database based on the mm10 mouse genome. Estimated counts and TPM values for transcripts were imported into R for downstream analysis. Prior to further analysis the expression and count values for transcripts were summed into gene level values. Differential expression testing was performed with DESeq2 using default options. Genes were marked as significantly differentially expressed with post-hoc adjusted p-values less than 0.05. <u>Gene Ontolgoy:</u> Functional annotation enrichment was performed for significantly mis- regulated genes against the background of all expressed genes in the samples using the hypergeometric test. Significantly enriched terms were clustered by constructing a nearest neighbors graph where edges were defined by shared genes associated with the terms being clustered. Terms that share more genes are considered “closer” to one another in the clustering compared to terms that share fewer or no genes at all. We used 10 nearest neighbors in the graph construction. The resulting networks were clustered using the igraph R package’s community detection function ‘cluster_infomap’ with default options.

### Statistical analysis

Except for RNA-sequencing analysis, all statistical analysis were performed using GraphPad Prism version 9.4.1. Appropriate tests are described in the corresponding figure legends. Significance was assigned according to Graphpad as: **ns** = p ≥ 0.05, **(*)** = 0.01 ≤ p < 0.05, **(**)** = 0.001 ≤ p < 0.01, **(***)** = 0.0001 ≤ p < 0.001, **(****)** = p < 0.0001.

## Acknowledgements

BTC core with funding from a NINDS Neuroscience Core Grant. The Peptide Synthesis Core Facility (Salk) with funding from NIH-NCI CCSG: P30 014195; the NGS Core Facility (Salk) with funding from NIH-NCI CCSG: P30 014195, the Chapman Foundation and the Helmsley Charitable Trust. The Transgenic Core Facility of the Salk Institute with funding from NIH-NCI CCSG: P30 014195. (UCSD-CMM-EM Core, RRID:SCR_022039 is supported in part by the National Institutes of Health Award number S10OD023527. The UCSD IGM Genomics Center with funding from a National Institutes of Health SIG grant (#S10 OD026929). ARD (Salk) for antibody production and animal care.

---

**Supplemental Figure 1.**
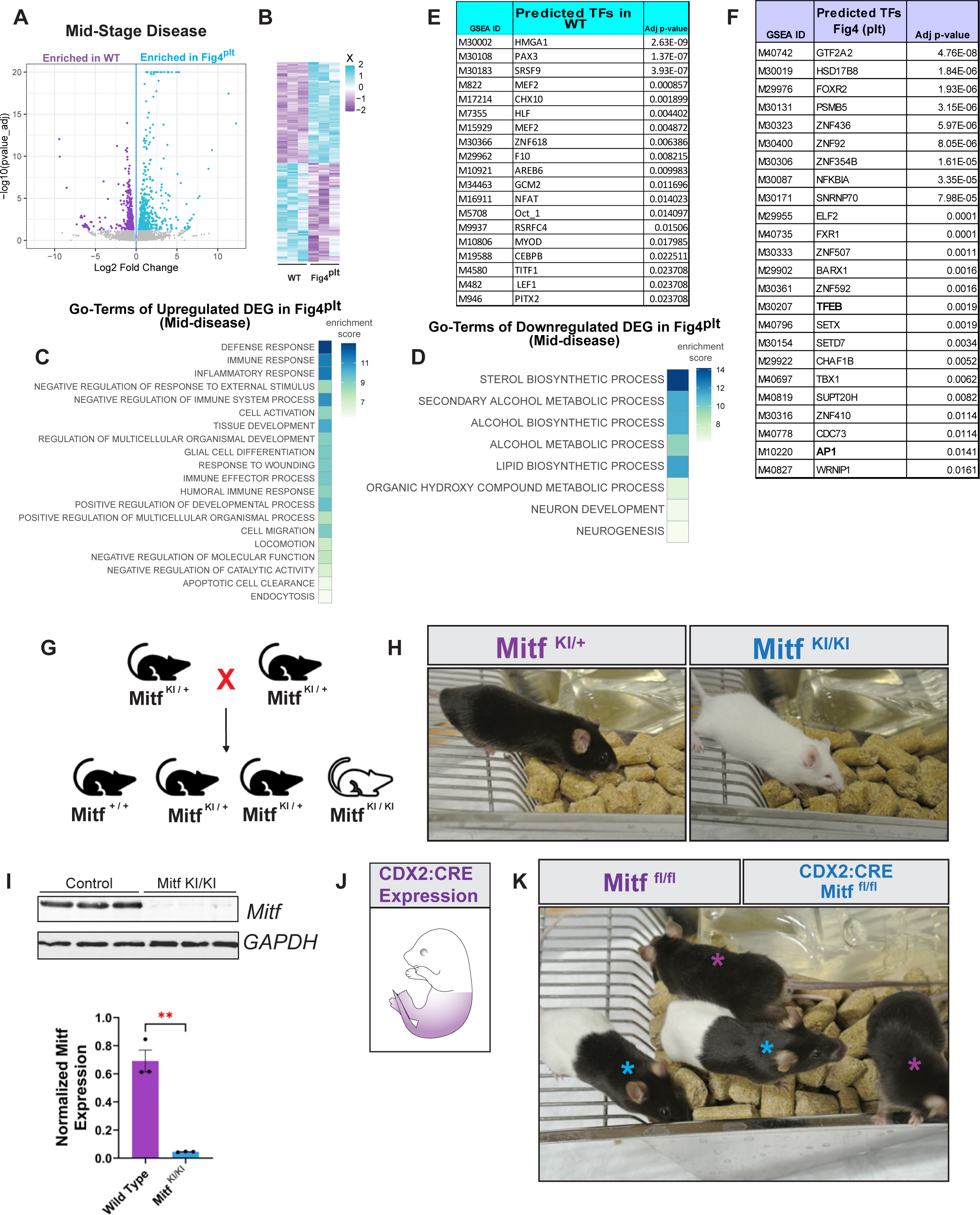
Network Analysis of Fig4^plt^ and Development of Mitf Conditional Allele. **(A)** Volcano plot showing log fold change versus adjusted p-value of sciatic nerves of Fig4^plt^ at postnatal day 12. Each dot represents a gene. Blue and purple dots are differentially expressed. (N= 3, each sample pooled from both sciatic nerves of 3 animals). **(B)** Heat map of expression levels (Z-score) of differentially expressed genes from **(A)**. Each row corresponds to a gene. **(C-D)** Gene Ontology (GO) term analysis using differentially expressed genes (DEG) as input compared with only those genes that are expressed in this experiment. The top two Biological Process (BP) GO-Terms from each cluster is shown. **(E-F)** Gene Set Enrichment Analysis (GSEA) of putative transcription factor binding enrichment in either WT **(E)** or Fig4^plt^ (P12) **(F)**. **(G)** Crossing scheme to generate homozygous Mitf^KI/KI^ animals. **(H)** Heterozygote animals for Mitf^KI^ allele exhibit normal pigmentation and eye size. Homozygous Mitf^KI/KI^ animals are albino with micropthalmia. **(I)** Sciatic nerve protein lysates from either control or Mitf^KO^ animals were subjected to analysis by SDS-page and immunoblotting for Mitf and GAPDH with the accompanying quantification of Mitf protein demonstrating a significant decrease in the level of Mitf protein expression of the Mitf^KO^ relative to the control (N=3). Unpaired t-Test. **(J)** Schematic of embryonic CDX2:CRE expression **(K)** The post germline FLP construct (Mitf^fl^) results in a functional conditional Mitf allele. Animals that are crossed to CDX2:CRE animals result in albinism in the caudal region of the animal concomitant with CDX2:CRE expression.

**Supplemental Figure 2.**
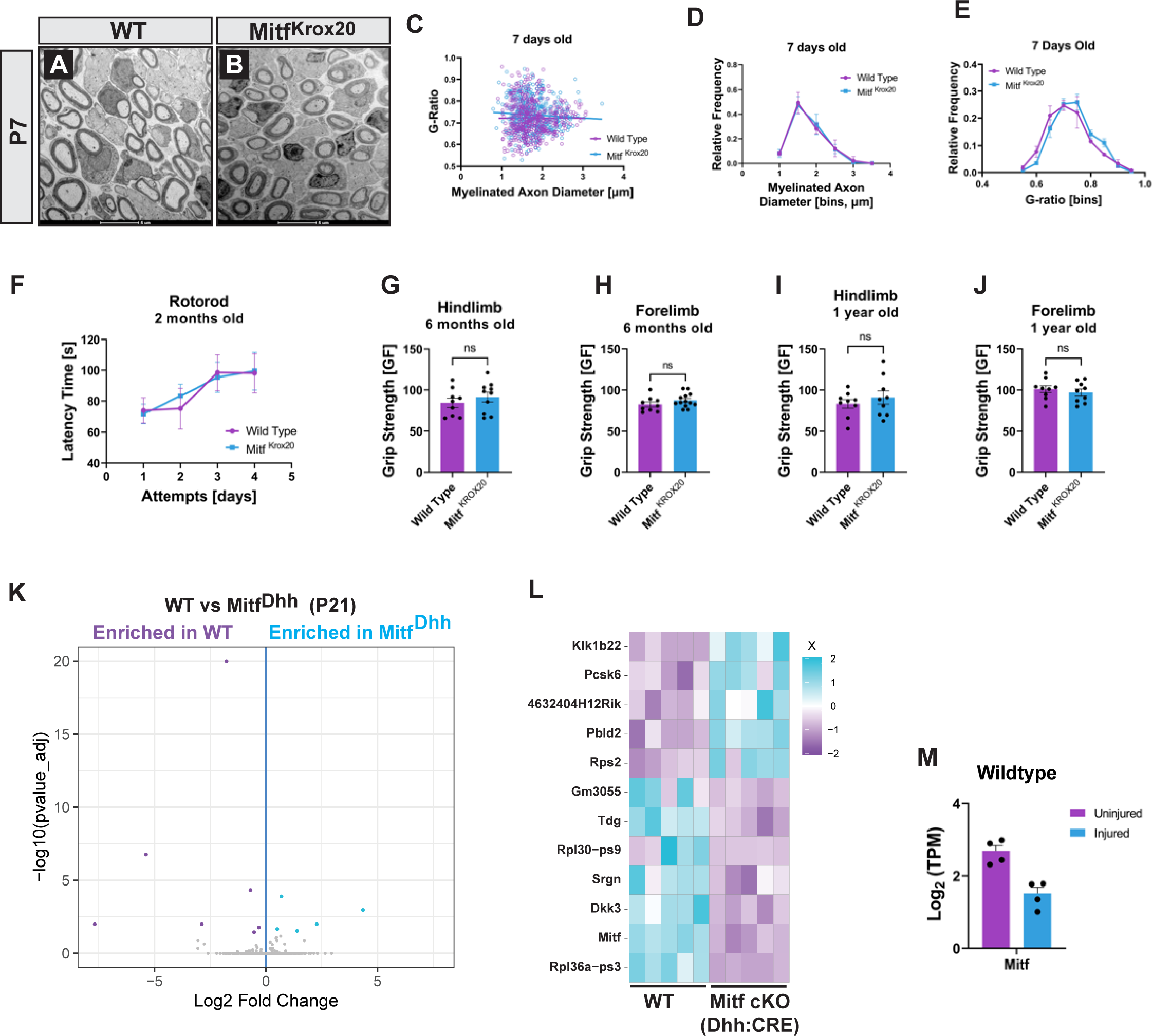
Developmental Analysis of Mitf Conditional Knockouts. (A-B) Representative TEM images of WT (A) and Mitf^Krox20^ (B) at P7. Scale bar = 5µM. **(C)** G-ratio plotted against axon diameter of P7 control (purple) and Mitf^krox20^ (blue) (n=3, at least 150 axons counted per sample). Lines represent simple linear regression. **(D)** Frequency of axon diameter in either WT or Mitf^Krox20^. Multiple unpaired t-Test (ns). **(E)** G-ratio frequency of WT and Mitf^Krox20^ (N=3, at least 200 axons counted per sample). Multiple unpaired t-Test (ns). **(F)** Rotorod of 2 month old WT (purple) or Mitf^Krox20^ (blue) shows no difference between the genotypes. Each day is an average of 4 trials per animal (n=8). Mixed effects ANOVA analysis with Geisser-Greenhouse’s correction: column factor (WT v Mitf^Krox20^) p = 0.9365 (ns), F (1, 14) = 0.006571; row factor (time) p = 0.0005 (**) F (1.747, 24.46) = 11.55; column x row factor p = 0.7161 (ns), F (3, 42) = 0.4537. **(G-H)** Hindlimb grip strength analysis at 6 months old between WT (N=9) and Mitf^Krox20^ (n=10) show no difference between genotypes **(G)**. Forelimb grip strength analysis at 6 months old between WT (n=9) and Mitf^Krox20^ (n=13) show no difference between genotypes **(H)**. Unpaired t-Test. **(I-J)** Hindlimb **(I)** and Forelimb **(J)** grip strength of 1 year old WT and Mitf ^Krox20^ animals (n=9) show no difference between genotypes. Unpaired t-Test. **(K)** Volcano plot showing log fold change versus adjusted p-value of sciatic nerve of WT or Mitf^Dhh^ at P21. Each dot represents a gene. Blue and purple dots are differentially expressed (n=5, each sample is pooled from both sciatic nerves of 3 animals). **(L)** Heat map of relative expression levels (Z-score) of differentially expressed genes from **(K)**. Each row corresponds to a gene. **(M)** Transcripts per million (TPM) of Mitf in uninjured sciatic nerves and in nerves 3 days post injury. Value of pAdj = ns.

**Supplemental Figure 3.**
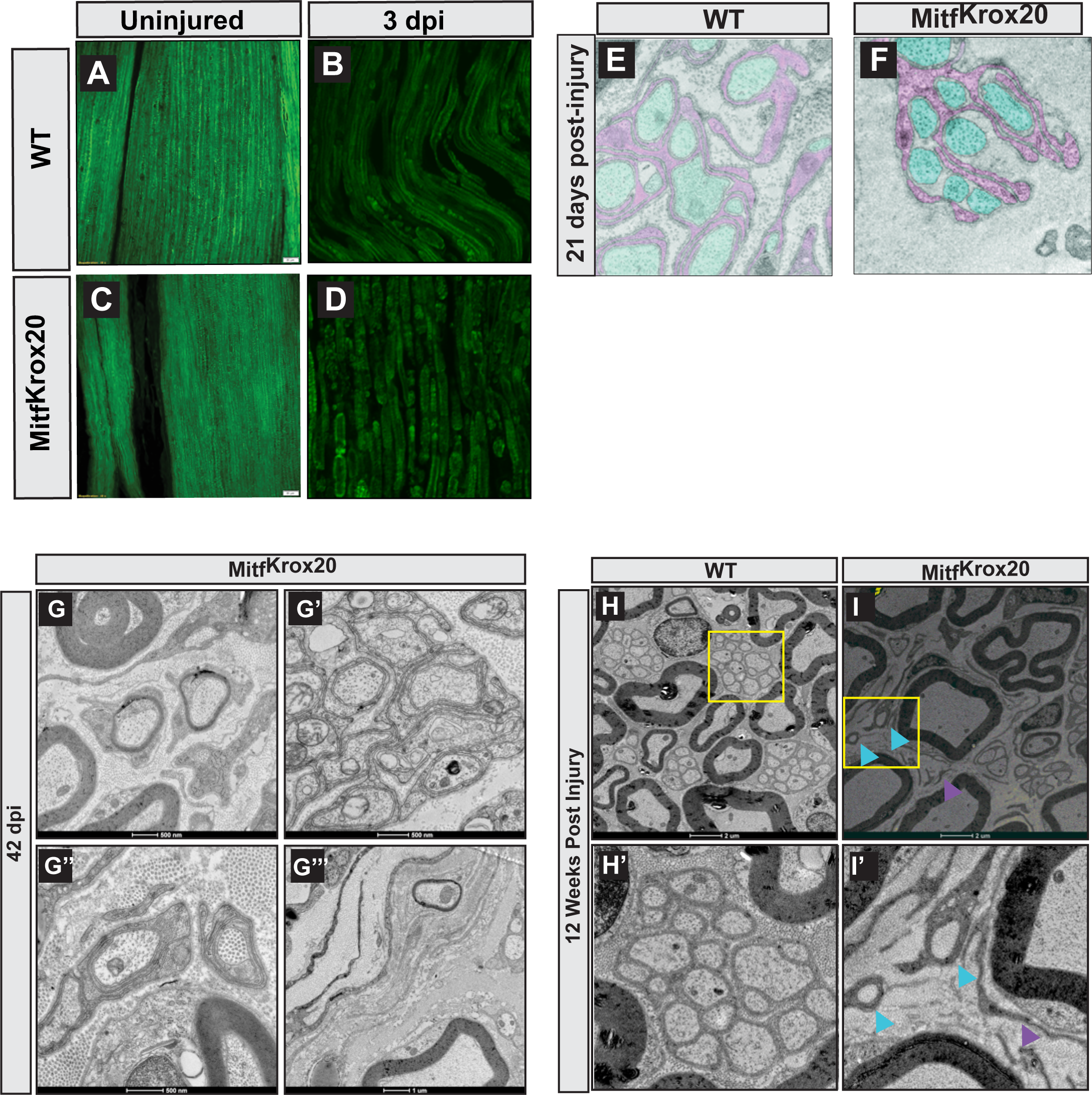
Time course of Mitf^Krox20^ After Injury. **(A-D)** Fluoromyelin-488 stained longitudinal sections of sciatic nerve of 8 weeks old animals in either uninjured or 3 dpi of WT or Mitf^Krox20^ animals. **(E-F)** Remak Schwann cell processes (pink) ensheath small caliber axons (blue) in the WT **(E)**. Remak Schwann cells in Mitf^Krox20^ show defects in establishing axonal contact **(F)**. **(G-G’’’)** TEM images from Mitf^Krox20^ showing ectopic myelination of small caliber axons **(G)**, supernumerated Schwann cell wraps around small caliber axons **(G’)**, Schwann cells wrapping around collagen in the extracellular matrix **(G’’)** and small caliber axon that is both ectopically myelinated and misprojecting through the perineurium **(G’’’)**. **(H-H’)** TEM images of the sciatic nerve from WT animals 12 weeks after sciatic nerve crush show normal recovery of Remak bundles. **(I-I’)** TEM images of the sciatic nerve from Mitf^Krox20^ animals, 12 weeks after sciatic nerve crush show the continued wrapping of collagen by Schwann cells (blue arrowheads). Schwann cell processes that do not contact axons (purple arrowhead).

**Supplemental Figure 4.**
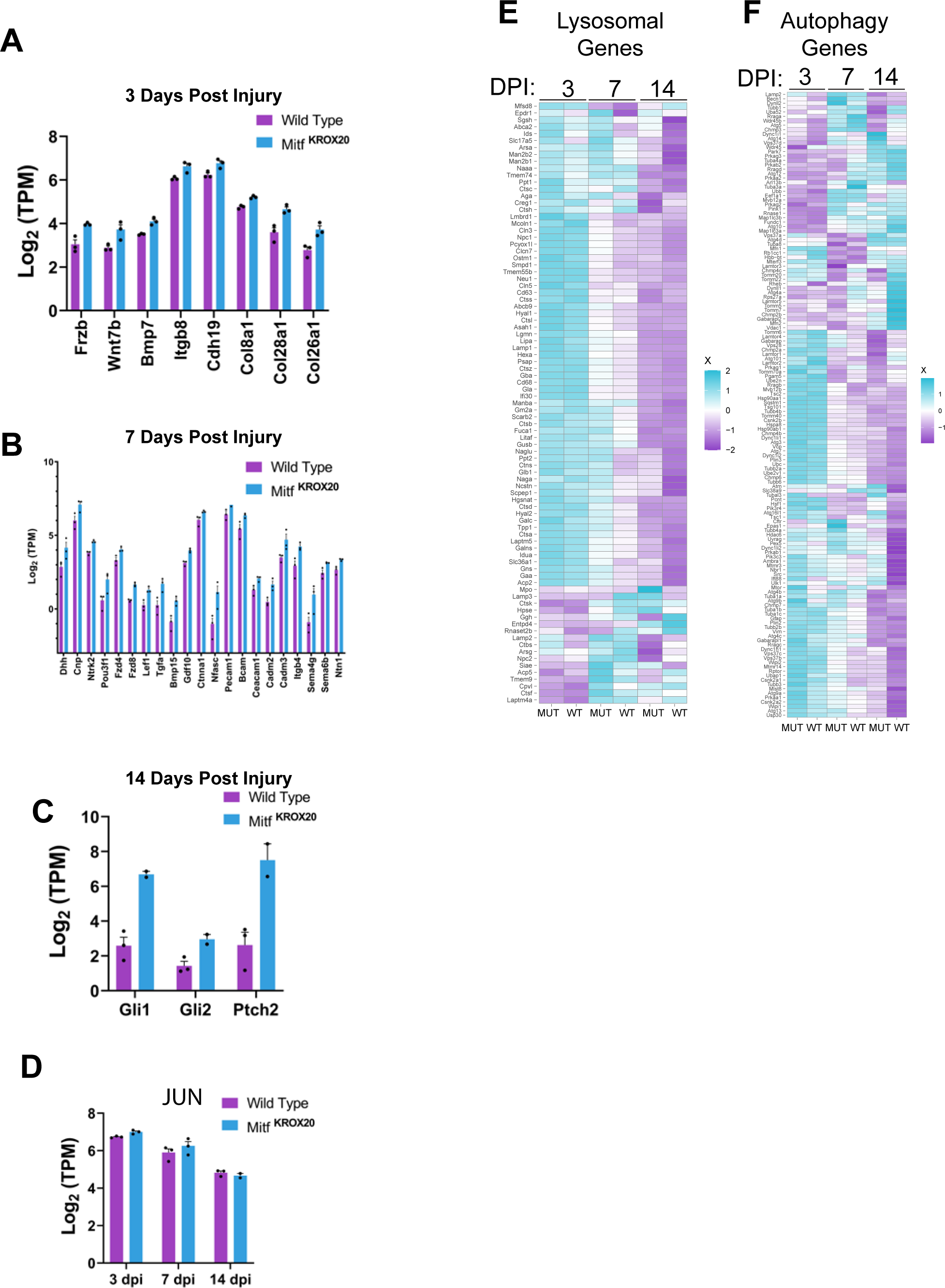
Gene Expression Analysis of *Mitf* mutants after Injury. **(A-C)** TPM (Transcripts per million) of selected differentially expressed genes from 3, 7 or 14 dpi in WT (purple) or Mitf mutants (blue). **(D)** TPM (Transcripts per million) of JUN at 3, 7 and 14 dpi in WT (purple) or Mitf mutants (blue). Value of pAdj = ns for all dpi. **(E)** Genes that constitute “Lysosome” Biological Process GO-term and their average Z-score expression in WT and Mitf mutant animals at 3, 7, and 14 dpi (Biological Process). **(F)** Genes that constitute “Autophagy” Biological Process GO-term and their average Z-score expression in WT and Mitf mutant animals at 3, 7, and 14 dpi (Biological Process).

**Supplemental Figure 5.**
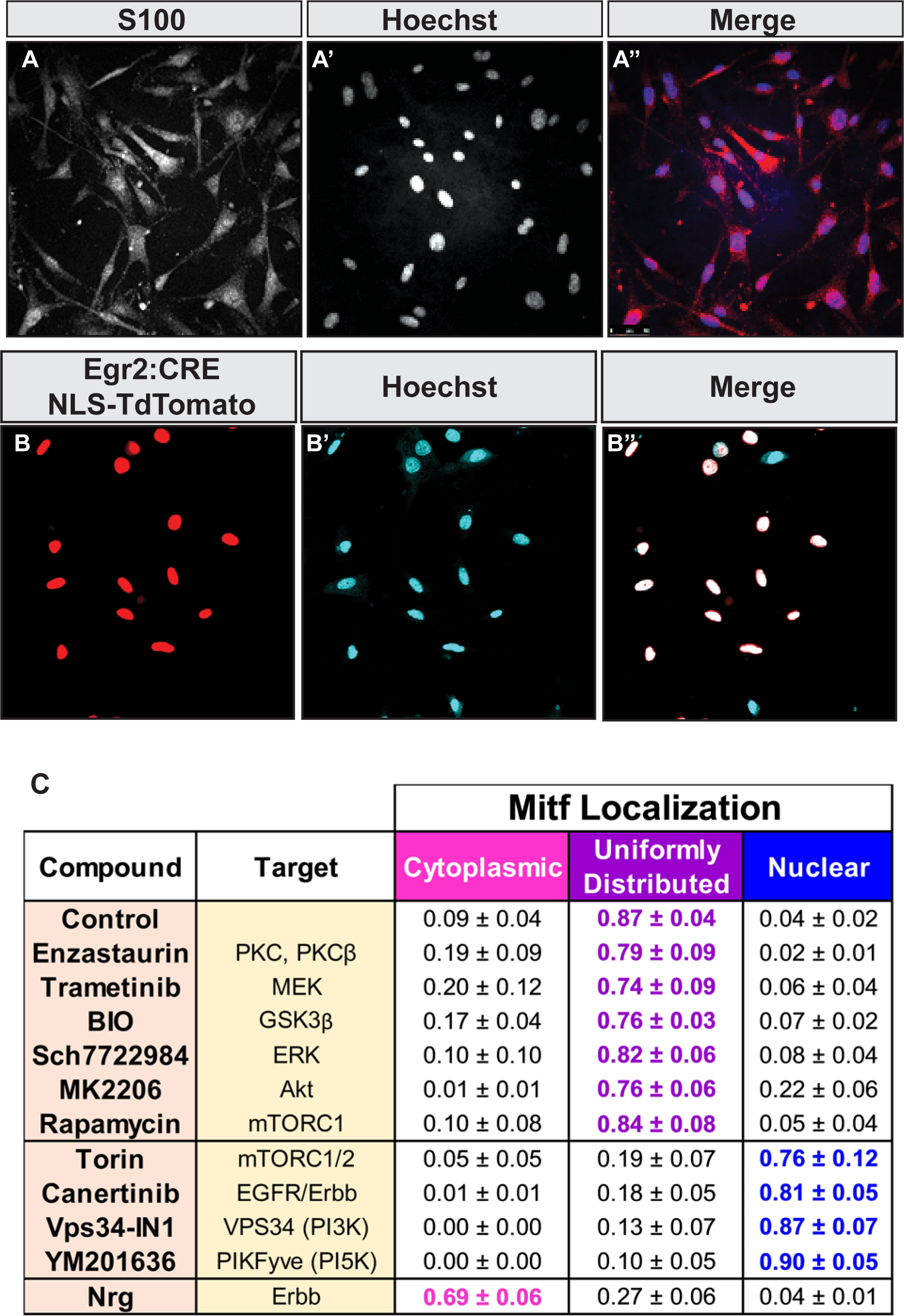
Primary Schwann Cell Culture. **(A-A’’)** Immunofluorescence of S100 in primary Schwann cell culture co-stained with Hoechst revealed that >97% of cells were labelled with S100 (N=3). **(B-B’’)** Schwann cell primary culture of genetically labelled Schwann cells using the Krox20 -CRE with a nuclear localizing Td-tomato (Ai75) reveal Schwann cells in culture. **(C)** Quantification of immunofluorescence of Mitf in primary Schwann cell culture. Mitf staining was uniformly distributed between the cytoplasm and nucleus in: control, Enzastaurin, Trametinib, BIO, Sch7722984, MK2206, Rapamycin conditions. Mitf was predominately nuclear in Torin, Canertinib, Vps34-IN1, YM201636. Mitf was predominately cytoplasmic after the addition of Nrg.

**Supplemental Figure 6.**
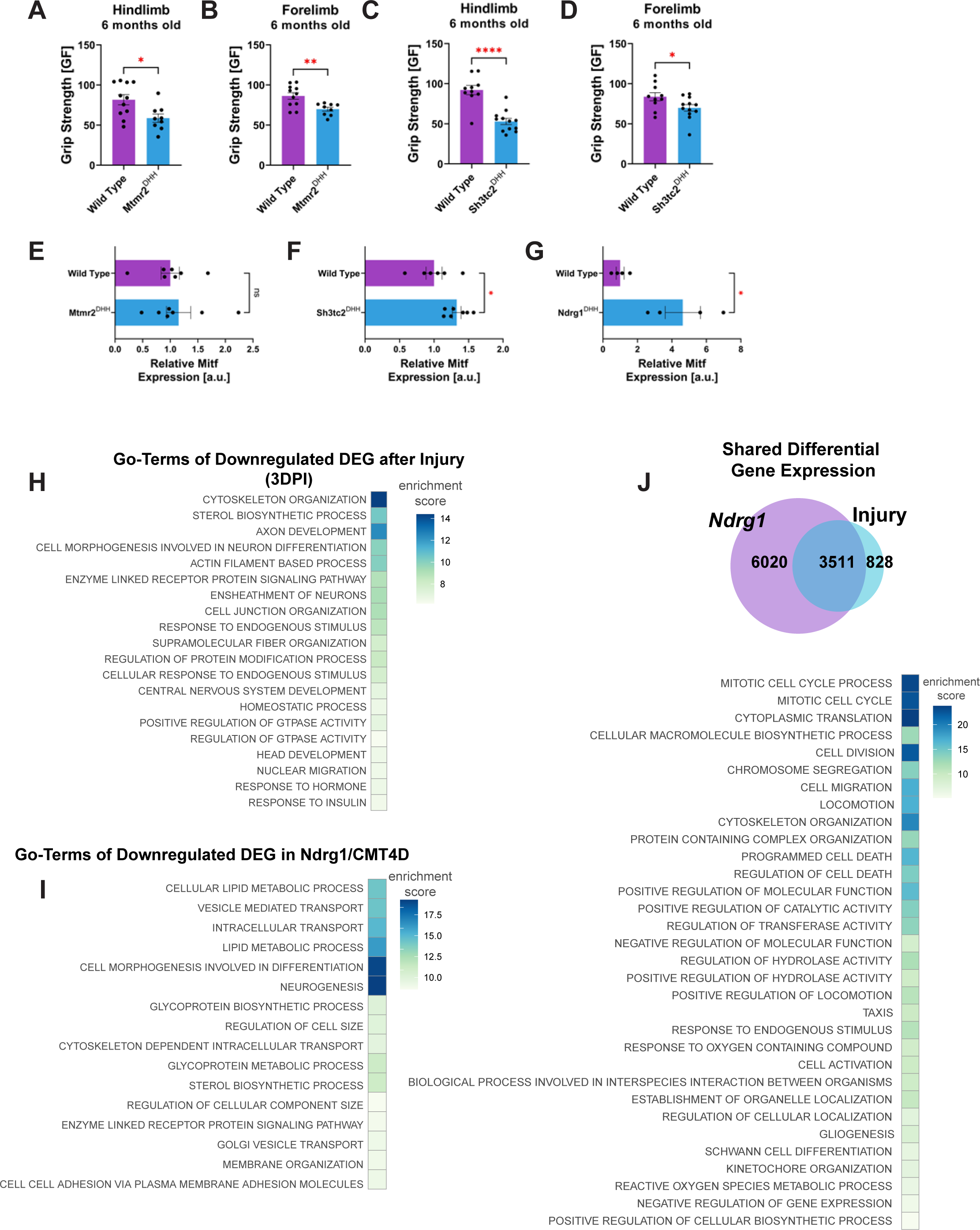
**(A)** Hindlimb grip strength of WT (n=11) and Mtmr2^Dhh^ (n=9) at 6 months old. Unpaired t-Test. **(B)** Forelimb grip strength of WT (n=11) and Mtmr2^Dhh^ (n=9) at 6 months old. Unpaired t-Test. **(C)** Hindlimb grip strength of WT (n=11) and Sh3tc2^Dhh^ (n=9) at 6 months old. Unpaired t-Test. **(D)** Forelimb grip strength of WT (n=11) and Sh3tc2^Dhh^ (n=9) at 6 months old. Unpaired t-Test. **(E-G)** Quantification of Mitf protein expression normalized against GAPDH and measured relative to WT (n=4-7) Each black dot represents an animal of the noted genotype. Unpaired t-Test. **(H)** Go-Term analysis of genes differentially downregulated after injury (3 days post injury) compared to uninjured. The top two Biological Process (BP) GO-Terms from each cluster are shown. **(I)** Go-Term analysis of genes differentially downregulated in Ndrg1^Dhh^ relative to WT at 4 weeks old. The top two Biological Process (BP) GO-Terms from each cluster are shown. **(J)** Venn Diagram depicting the shared differential expression between Ndrg1^Dhh^, and injured nerves (3dpi); 3511 genes are differentially expressed in both categories. Heat map of Go-Terms (Biological Process) from shared differentially expressed genes in both Injury and Ndrg1^Dhh^. The top two Biological Process (BP) GO-Terms from each cluster are shown.

**Supplemental Figure 7.**
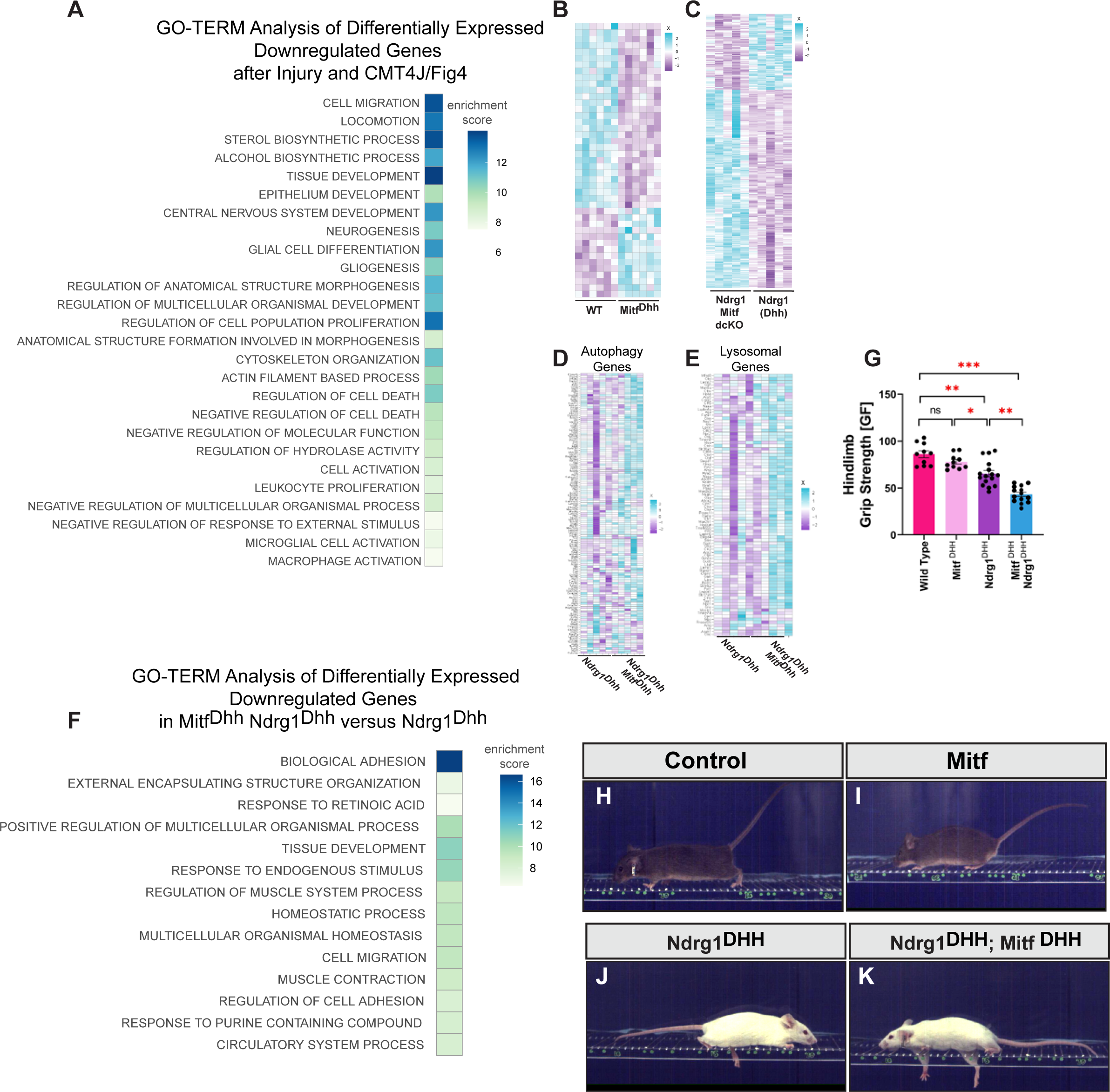
**(A)** Heat map of Go-Terms (Biological Process) from genes that are differentially expressed and downregulated in both Injury and Fig4^Plt^ compared to their respective controls. **(B)** Heatmap of expression levels (in Z-scores) of differentially expressed genes between WT and Mitf^Dhh^ at 4 weeks old. Each row corresponds to a gene. **(C)** Heatmap of expression levels (in Z-scores) of differentially expressed genes between Ndrg1^Dhh^ and Ndrg1^Dhh^; Mitf^Dhh^ at 4 weeks old. Each row corresponds to a gene. **(D)** Genes that constitute Autophagy GO-terms and their average Z-score expression in Ndrg1^Dhh^ and Ndrg1^Dhh^; Mitf^Dhh^ animals (Biological Process). **(E)** Genes that constitute Autophagy GO-terms and their average Z-score expression in Ndrg1^Dhh^ and Ndrg1^Dhh^; Mitf^Dhh^ (Biological Process). **(F)** Heat Map of GO-Terms (Biological Process) of differentially expressed and downregulated genes in Ndrg1^Dhh^; Mitf1^Dhh^ compared to Ndrg1^Dhh^. **(G)** Hindlimb grip strength of WT (n=10), Mitf^Dhh^ (n=10), Ndrg1^Dhh^ (n=16) and the double conditional knockout Ndrg1^Dhh^Mitf ^Dhh^ (n=14) animals at 8 weeks. Ordinary One-way ANOVA with Tukey’s multiple comparison test. **(H-K)** Images of wildtype **(H)**, Mitf^Dhh^ **(I)**, Ndrg^Dhh^**(J)** and Ndrg1^Dhh^; Mitf^Dhh^ **(K)** walking across the irregular ladder beam.

## Notes

### Competing Interest Statement

The authors have declared no competing interest.

